# Disruption of *OsLAT5* is sufficient to endow rice tolerance to dihydropyridine herbicides at commercial application concentrations

**DOI:** 10.1101/2022.11.06.515312

**Authors:** Ronghua Chen, Di Zhao, Diya Yu, Chaozheng Li, Siwei Wang, Hanhong Xu, Fei Lin

## Abstract

Breeding non-selective herbicide-resistant crops is important constituent of weed management system in modern rice production. Non-selective dihydropyridine herbicide shares transporters with polyamine (PA), making construction of a dihydropyridine herbicide-resistant rice line possible by inactivating the PA transporter function via gene editing. Success depends on understanding substrate selection for homologues in the PA transporter family and amino acid sites that play critical roles. Here, *OsLAT1* was mainly responsible for root uptake and root-to-shoot transport; whereas, *OsLAT5* was more responsible for intracellular transport to chloroplasts. The *oslat5* disruption line tolerated relevant concentrations, while *oslat1* did not. Compared to GY11 wild type, plant height, 1000-grain weight, and spermidine, spermine, and putrescine content changes occurred in *GY11-oslat5* lines, implying involvement of *OsLAT5* in yield and quality regulation. *OsLAT5_P44F, P44Y and P44R_* showed declined dihydropyridine herbicide uptake but no spermidine and putrescine transport competence disruption in yeast, providing a candidate site for precisely editing in breeding a dihydropyridine herbicide-resistant rice cultivar without impairing rice yield and grain quality.

## Main text

Rice direct seeding and other simplified cultivation methods represent the direction of the development of rice culturing, among which proper weed control is of great importance to maximize yields and stability^1,2^. Coordination of herbicide applications with herbicide-resistant crops is a favorable strategy for controlling weeds in paddy fields. Non-selective herbicides are widely used due to the simplicity of application and low cost. However, these herbicides kill indiscriminately, affecting both weeds and crops^3^. Thus, it is necessary and of great significance for simplifying rice cultivation to cultivate non-selective herbicide-resistant rice. To date, only a few non-selective herbicide-resistant rice cultivars have been reported and none have been commercialized.

Dihydropyridine herbicides belong to an important and widely used category of non-selective herbicides, including paraquat (1,1-dimethyl-4,4-bipyridinium dichloride) and diquat (1,10-ethylene-2,20-bipyridinium dibromide)^4^, with the characteristics of high contact-killing efficiency^5^, limited long-distance transport activity affected by illumination^6,7^, and biological inactivity in soil^8–10^. After application, dihydropyridine herbicides rapidly kill green plant tissue under light conditions by harvesting electrons from photosystem I (PSI) and transferring them to molecular oxygen, thus stimulating the overproduction of reactive oxygen species (ROS), which can cause membrane damage^11–13^. The action mode of dihydropyridine herbicides means that they reach the target site from extracellular space via the uptake and transport process. As early as 1974, paraquat accumulation in rat lung was confirmed as an energy-dependent process that obeyed saturation kinetics^14^, implying that paraquat uptake is mediated by transporters. In plants, an uptake study was first performed in maize root in 1992 and indicated that paraquat was transported into cells by a protein-mediated saturable system^15^. Other studies revealed that a series of polyamines (PAs) competitively inhibited paraquat transport and thereby enhanced plant resistance which counteracted paraquat toxicity^16–20^. Paraquat meets the structural criteria of being PA-like, with at least two charged nitrogen atoms and a polar structure, and so is allowed transportation into cells via the PA transport system^13,21,22^. Knowledge of paraquat transporters in plant largely comes from the model plant *Arabidopsis thaliana*. RMV1 (resistant to methyl viologen1) was first identified as functioning in PA and paraquat uptake, which encodes a L-type amino acid (LAT) transporter^23,24^. Another LAT transporter par1 (paraquat resistant1) is involved in intracellular transport of paraquat into chloroplasts^25^, and then the mutant *par2* generated from nonsense mutation of PAR1 affects transportation of putrescine^26^. PDR11 belonging to an ATP-binding cassette (ABC) transporter was shown to be responsible for paraquat accumulation in plant cells and could be inhibited by putrescine^27^. Another ABC transporter B family protein, PqTS2, was identified in weed goosegrass^28^; it is a homolog of ABCB1 in human and mice involved in paraquat efflux^29^, and there appears to be a relationship between PA transport and paraquat resistance. Recently, the multidrug and toxic extrusion (MATE) family transporter DTX6 was shown to confer paraquat resistance by exporting paraquat out of the cytosol, which differed from the above-mentioned PA transport system^30,31^. Overall, it is evident that paraquat enters plant cells via various transport systems.

The LAT family is one of the main plasma membrane transporters that likely play considerably important roles in paraquat transport in plants. LATs are responsible for the permeation of the amino acid-PA-and organocations through the plasma membrane^32–34^, and studies have shown that LAT1 also mediated the transport of amino acid-related drugs^35,36^, suggesting broad substrate selectivity of LATs. In the model plant species *Arabidopsis thaliana*, there are five transporters (LAT1–5)^24,34^, two of which (AtLAT1/RMV1/AtPUT3 and AtLAT4/PAR1/AtPUT2) exhibited paraquat and PA transport activity^23,25^. Rice consists of eight gene-encoding LAT proteins, in which several LATs have been identified. For example, OsLAT1/OsPUT1, OsLAT5/OsPUT3, and OsLAT7/OsPUT2 were involved in paraquat and PA transport as confirmed by studies on yeast^37,38^. Furthermore, knocking_down *OsLAT5/OsPAR1* expression in rice by an RNA interference (RNAi) construct exhibited paraquat resistance^25^. In addition, the mutagenesis *OsLAT1/5/7* improves paraquat tolerance in rice^39^.

The concept of construction of herbicide-resistant germplasms by blocking transporters is promising. Disruption of herbicide transporters will decrease herbicide uptake and accumulation in cells, which not only makes the plants more herbicide resistant but also reduces the risk of residual herbicides in plants. CRISPR-mediated gene editing technologies have merits of being highly efficient and precise, which make it possible to develop crops with herbicide transportation deficiency without being compromised in economic traits such as yield, eating quality, and disease resistance, etc.^40,41^. Therefore, extensive study of dihydropyridine herbicide transporters in rice, and characterization of their affinity to the substrate and to both diquat and paraquat, would provide insight into breeding a dihydropyridine herbicide-resistant rice germplasm with commercial value. In this study, we aimed to dissect the functionally differentiation between *OsLAT1* and *OsLAT5* in dihydropyridine herbicide transport in yeast and rice plants. A variety (in the Guiyu No. 11 background) with the *OsLAT5* mutation was generated and demonstrated the commercial application value on the rice cultivation under controlled and field conditions.

## Results

### *OsLAT1* and *OsLAT5* show dihydropyridine herbicide transport activity in yeast

Phylogenetic relationships revealed that PA transporter domains were highly conserved between *A. thaliana* and *Oryza sativa. OsLAT1* and *OsLAT5* clustered in a clade with *AtLAT1* and *AtLAT4*, the two homologous that exhibited PA transportation ability^38^(Fig. S1A), suggesting they may have been tightly involved in dihydropyridine herbicide transport. To explore the relationships among rice *LAT* genes and dihydropyridine herbicide transport, we selected four genes for analysis: *OsLAT1* (LOC_Os02g47210), *OsLAT5* (LOC_Os03g25869), *OsLAT3* (LOC_Os03g37984), and *OsLAT7* (LOC_Os12g39080). *OsLAT3* and *OsLAT7* were selected as they are formed in the other clade and could be used as negative controls. Given that the PA transporter was able to identify dihydropyridine herbicide with structural similarity, the Δ*agp2* that was a PA uptake-deficient mutant of yeast was also used to screen genes involved in transport of these herbicides^42^.

The Δ*agp2* strain carrying empty vector (pYES-dest52) lacked the activity to uptake dihydropyridine herbicide; therefore, it could survive on SD-Gal medium containing this herbicide; whereas, growth of the BY4741-empty vector (WT) strain was inhibited. These four genes were expressed under the GAL1 promoter in Δ*agp2* strain. Compared to *OsLAT3* and *OsLAT7*, the yeast transformed with *OsLAT1* and *OsLAT5* complemented the Δ*agp2* phenotype and conferred sensitivity to 0.75 mM of both paraquat and diquat. The result clearly showed transport competence of *OsLAT1* and *OsLAT5*, but not of *OsLAT3* and *OsLAT7* (Fig. S1B). The yeast growth and uptake assay were conducted for *OsLAT1* and *OsLAT5*. The yeast harboring *OsLAT1* or *OsLAT5* became sensitive with an increasing dihydropyridine herbicide concentration, and *OsLAT5* showed higher sensitivity (Fig. 1A). The growth curves also agreed with the results observed on solid medium (Fig. 1B-E). The uptake exhibited Michaelis–Menten saturation kinetics, with *K_m_* rankings of *_KmOsLAT1_* > *_KmOsLAT5_* for both herbicides (Fig. 1F–J and S2). The result provided evidence supporting the idea that *OsLAT1* and *OsLAT5* are PA and PA structural analogue—dihydropyridine herbicide transporters, with *OsLAT5* having stronger activity.

**Fig. 1.**
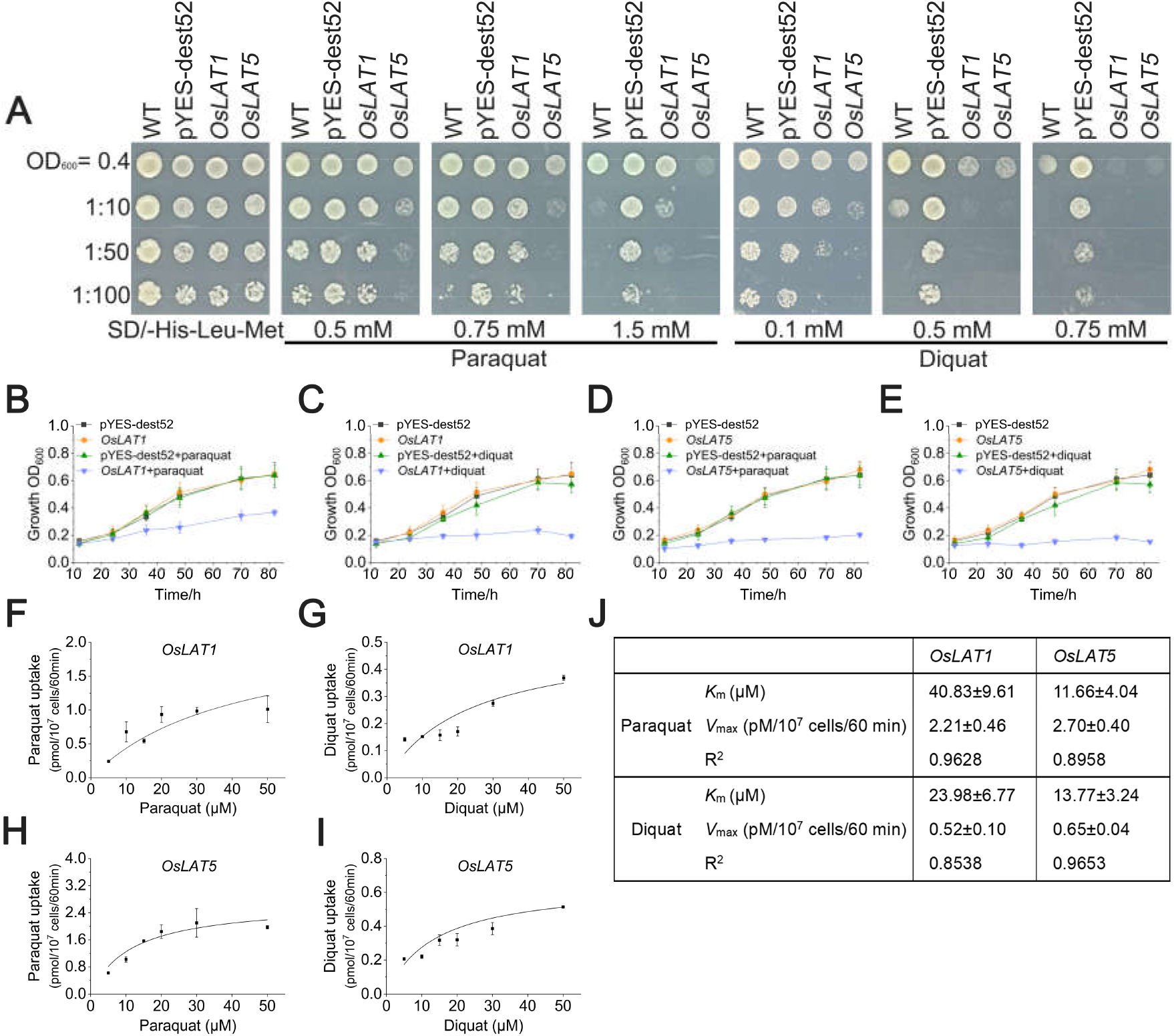
*OsLAT1* and *OsLAT5* function as dihydropyridine herbicide transporter in yeast heterologous system. (A) Growth phenotypes of BY4741 and Δ*agp2* on solid medium supplemented with dihydropyridine herbicide or polyamines at the indicated concentrations. BY4741 strain was transformed with empty vector (WT), and Δ*agp2* strains were transformed with empty vector (pYES-dest52) or pYES-dest52 carrying *OsLAT1* or *OsLAT5*. Plates were photographed after 5 days of incubation at 30°C. (B-E) Growth curves plotted from OD_600_ values of liquid cultures. Yeast cells from the strains described in (A) were grown in liquid medium containing 0.75 mM paraquat or diquat. Growth of cultures was monitored by taking OD_600_ readings in the indicated time (12, 24, 36, 48, 70 and 82 h). Data are mean ± SD. n = 3. (F-J) Dose-dependent and kinetic analysis of dihydropyridine herbicide uptake by *OsLAT1* and *OsLAT5*. Michealis–Menten curves for dihydropyridine herbicide uptake were obtained by subtracting the uptake in vector control expressing cells from that in Δ*agp2-OsLAT1* or *OsLAT5* cells.

### Transcription pattern of *OsLAT1* and *OsLAT5* and its subcellular localization

The real-time polymerase chain reaction (RT-PCR) results and examination of the RiceXPro microarray data showed that *OsLAT5* were expressed in all tissues at a relatively high level compared to *OsLAT1*. Furthermore, both *OsLAT1* and *OsLAT5* were mainly expressed in roots and embryos (Fig. S3A and 3B). Time-dependent RT-PCR analysis showed that *OsLAT1* and *OsLAT5* expressions were consistently induced by dihydropyridine herbicide exposure, with more significant up-regulation in *OsLAT1* (Fig. S3C). To assess subcellular localization, the OsLAT1-GFP or OsLAT5-GFP fusion protein was transiently co-expressed in *A. thaliana* protoplasts with various cell organelle markers. OsLAT1-GFP fluorescence was colocalized with the plasma membrane marker 1008s-mCherry, whereas, the OsLAT5-GFP fluorescence signal was observed in both the plasma membrane and Golgi apparatus (Fig. S3D).

Combining the results of the yeast assay and expression pattern analysis, we suggest that there are differences between *OsLAT1* and *OsLAT5* in transport activity and gene expression, implying that they perform different roles in dihydropyridine herbicide uptake and transportation in plants.

### Tolerance to dihydropyridine herbicide during germination mediated by *OsLAT1* and *OsLAT5*

To check the dihydropyridine herbicide-sensitive phenotype, we generated rice lines carrying various *OsLAT1* or *OsLAT5* genotypes (in Zhonghua11 background), two independent mutants edited by CRISPR-Cas9 (*OsLAT1-a* and *OsLAT1-b* or *oslat5-c* and *oslat5-d*) and two overexpression lines of *OsLAT1* and *OsLAT5* driven by a 35S promoter (*OsLAT1-OE8* and *OsLAT1*-OE10 or *OsLAT5-OE2* and *OsLAT5-OE9*) (Fig. S4).

The germination rate of different *OsLAT1* or *OsLAT5* genotype lines was not affected by the addition of 0.5 μM diquat. On day 6, the primary growth of *oslat5* mutants in medium containing 0.5 μM diquat was optimal among the tested rice lines (Fig. 2B). In the presence of 0.05 μM diquat, the *OsLAT1-OE* lines displayed phenotypes with severely decreased tolerance, with reduced shoot and root lengths compared to wild-type and *oslat1*mutants (Fig. S2C). As the diquat concentration increased to 0.1 μM, growth of *OsLAT1*genotype lines was almost inhibited completely (Fig. S2E). None of the *OsLAT5* lines revealed any difference in the absence of diquat, which was consistent with the result in *OsLAT1* lines. Under the assay conditions, *oslat5* mutants displayed excellent resistance to diquat even when the diquat concentration increased to 0.1 μM, whereas wild-type and *OsLAT5*-OE seedlings exhibited similar inhibited shoot and root growth (Fig. S2D and F).

**Fig. 2.**
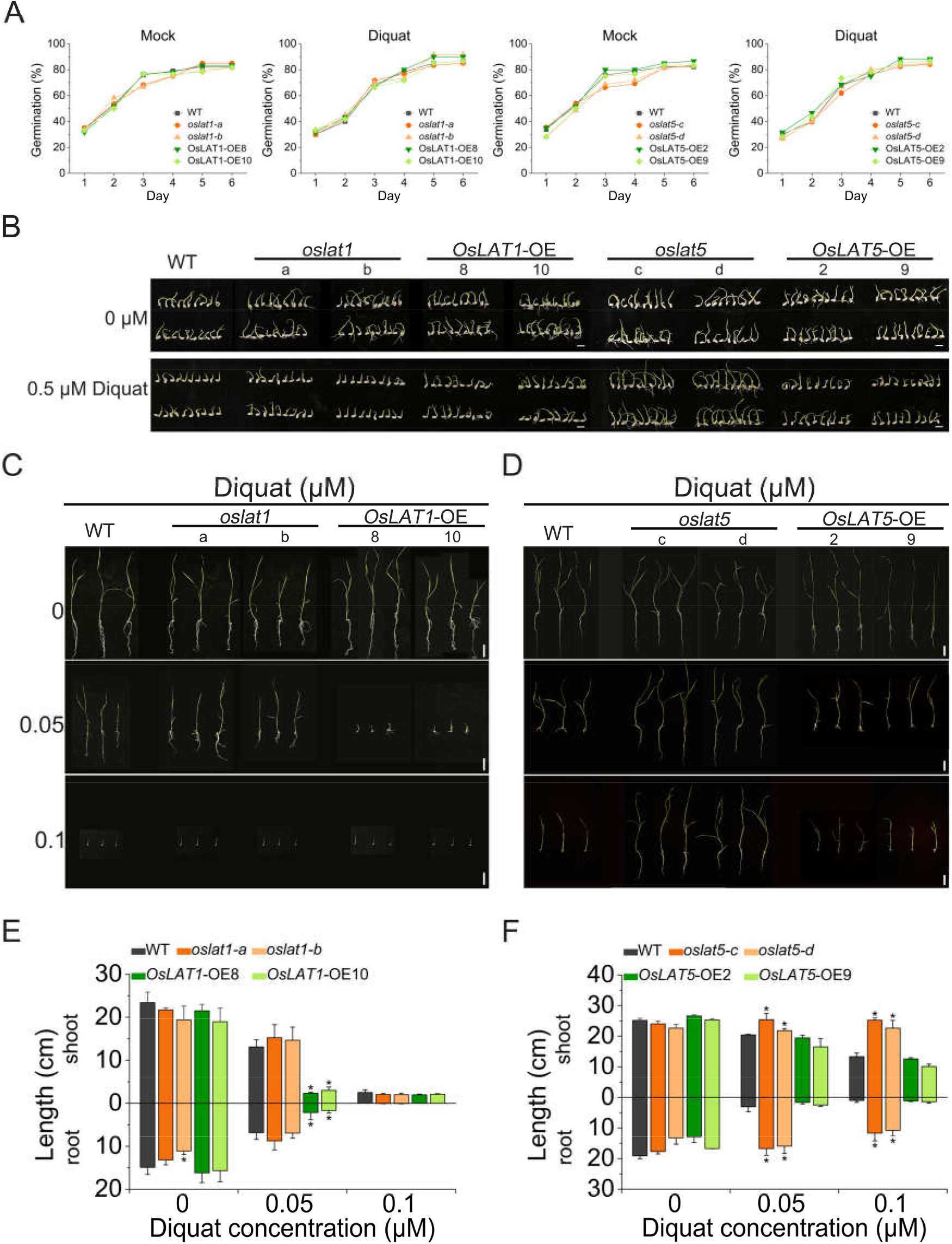
Rice seed germination and seedling growth on medium containing diquat. (A) Seed germination curve. Seeds of wild type, mutants and over-expression line of *OsLAT1* or *OsLAT5* were germinated on MS medium added with 0 or 0.5 μM diquat for 6 days and germination rate was recorded every day. (B) The above seeds continued to grow followed by six days growing before pictures taken. Bar = 1 cm. (C-D) Growth of germinated rice seedlings mediated by *OsLAT1* (C) or *OsLAT5* (D) with 0.05 and 0.1 μM diquat on MS medium. Bar = 4 cm. (E-F) The shoot and root length of rice seedlings shown in (C) and (D). Data are mean ± SD. n = 3. Asterisks indicate significant differences (one-way ANOVA: *\*P* < 0.05).

Similarly, *OsLAT1* or *OsLAT5* genotype lines exhibited consistent growth with exposure to paraquat and diquat (Fig. S5). These findings suggested that, in the seedling stage, overexpression of *OsLAT1* rendered rice plants sensitive to dihydropyridine herbicides; however, blocking of the *OsLAT1* function did not endow tolerance to the herbicides. Conversely, rice seedling deficiency of *OsLAT5* conferred dihydropyridine herbicide tolerance; however, its overexpression could not change such tolerance.

### Knockout of *OsLAT5* endowed rice tolerance to dihydropyridine herbicide at the tillering stage

To determine the effects of commercially available dihydropyridine herbicide on the growth and development of different *OsLAT1* and *OsLAT5* genotype rice lines, we tested 4-week-soil-grown rice that was foliar sprayed with 50, 100, or 200 mg/L paraquat or diquat under a 28-d observation. The Fv/Fm and chlorophyll content in plants exposed to herbicide were measured to assess the photosynthetic performance and degree of plant wilting.

When treated with 50 or 100 mg/L diquat, the growth and development status of wildtype, *oslat1* mutants, and *OsLAT1-OE* were equivalent. However, under the higher diquat concentration (200 mg/L), overexpression of *OsLAT1* conferred damage to the transgenic rice from the 8^th^ after spraying (Fig. 3A and B). Similar to the germination assay results, mutation of *OsLAT5* remarkably increased diquat tolerance, leading to survival of *oslat5* mutants during the observation period. In contrast, the *OsLAT5*-OE seedlings became hypersensitive, even when treated with relatively lower diquat concentration. Surprisingly, *oslat5* sprayed with 200 mg/L diquat showed sensitivity with some leaf injury; however, a remarkable recovery occurred over time, as shown by the Fv/Fm and chlorophyll content. For example, on the 28^th^ day, *oslat5* exhibited 10.8% (*oslat5-c*) and 14.2% (*oslat5-d*) reductions in Fv/Fm compared with 58.0% (WT) and almost 100% (*OsLAT5*-OE) reductions (Fig. 3C and 3D).

**Fig.3.**
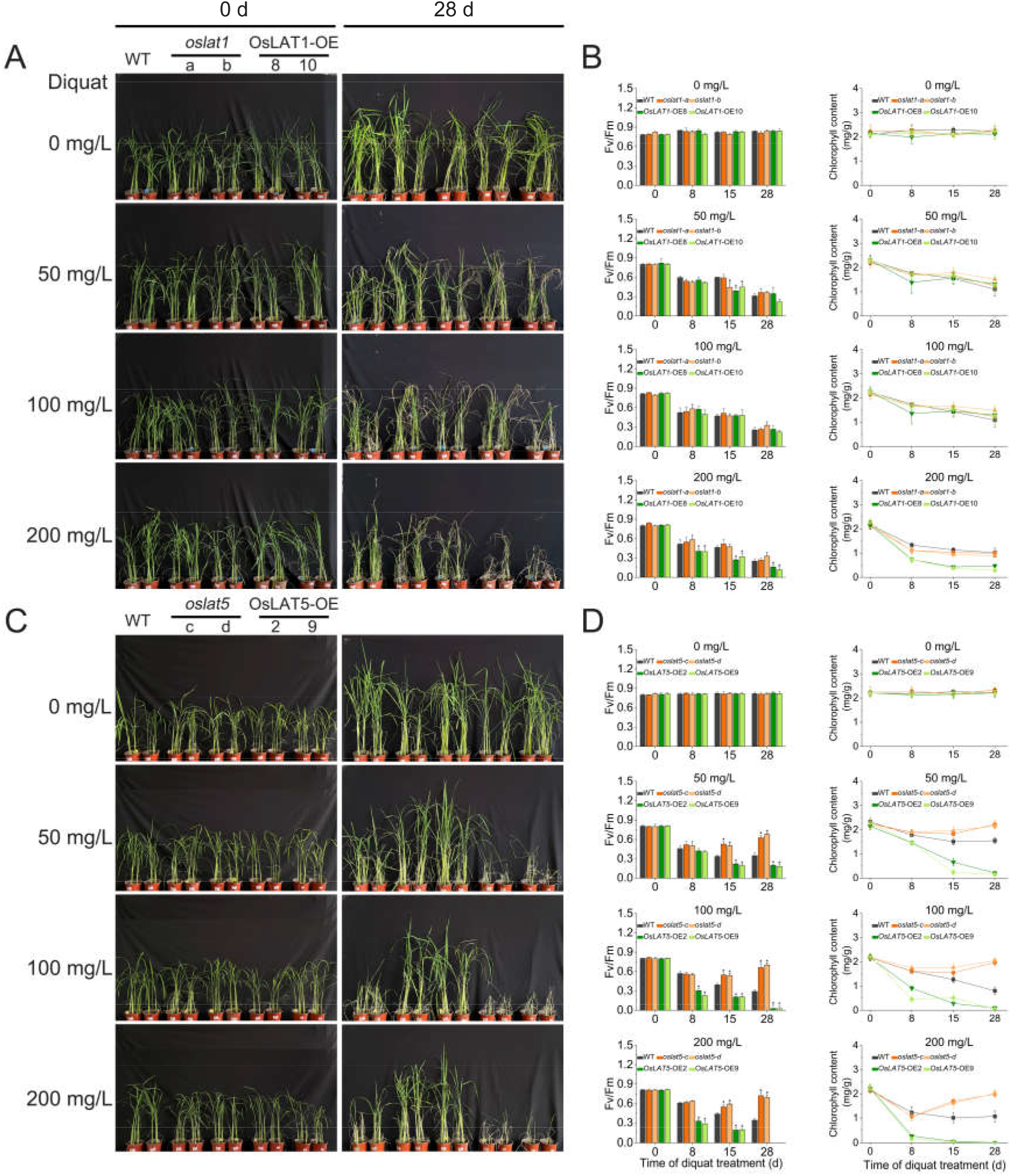
Effects of foliar spraying of diquat on rice growth and development at tillering stage. (A) Field-grown rice of different lines of *OsLAT1*(4 weeks old) sprayed with 50, 100 and 200 mg/L diquat followed by continue growth for different durations. (B) Analysis of Fv/Fm and chlorophyll contents in the leaves from (A) treated with diquat for the indicated durations. Data are mean ± SD. n = 3. Asterisks indicate significant differences (one-way ANOVA: *\*P* < 0.05). (C) Field-grown rice of different lines of *OsLAT5* (4 weeks old) sprayed with 50, 100 and 200 mg/L diquat followed by continue growth for different durations. (D) Analysis of Fv/Fm and chlorophyll contents in the leaves from (C) treated with diquat for the indicated durations. Data are mean ± SD. n = 3. Asterisks indicate significant differences (one-way ANOVA: \**P* < 0.05).

Paraquat had a detrimental effect on wild-type, *oslat1* mutants, and *OsLAT1*-OE, and all died within 28 days of application. Moreover, the Fv/Fm and chlorophyll contents decreased as the paraquat concentration increased, and there were no differences among *OsLAT1* lines (Fig. S6A and B). The seedlings of *oslat5* mutants showed a degree of resistance to paraquat especially under 50 mg/L and 100 mg/L treatment, while wild type and *OsLAT5*-OE suffered consistent phytotoxicity at all concentrations. We also measured the Fv/Fm and Chlorophyll contents, their reduction in *oslat5* mutants were less than in wild type and *OsLAT5*-OE (Fig. S6C-S6D). These findings indicated that *OsLAT5*enhanced rice dihydropyridine herbicide resistance at the commercial application concentration, which is of great significance for field applications. Although *OsLAT1* was involved in the transportation of dihydropyridine herbicide, its mutation did not confer resistance to such herbicides in rice plant.

### Uptake and accumulation of dihydropyridine herbicide in rice mediated by *OsLAT1*and *OsLAT5*

To further characterize the biochemical basis of *OsLAT1* and *OsLAT5–* dihydropyridine herbicide interaction, analyses of the root uptake and shoot translocation of dihydropyridine herbicide in various *OsLAT1* and *OsLAT5* genotypes rice were performed.

The total amount of diquat of *OsLAT1-OE* was markedly increased in the 5 and 10 μM concentrations; whereas, no obvious difference in diquat uptake was observed in wild type and *oslat1* mutants. We also measured the diquat content in rice root, stem, and leaf to further assess the transport activity of *OsLAT1*.During the diquat incubation, there was much more diquat uptake and transport in *OsLAT1-OE* than in wild type and *oslat1* mutants. However, the poor dihydropyridine herbicide translocation ability^43,44^, which resulted in undetectable contents in leaves, resulted in no differences among three *OsLAT1* rice genotypes (Fig. 4A). Time-dependent accumulation experiments showed that diquat accumulation plateaued in wild type, *oslat1* mutants, and *OsLAT1-OE* with no difference for the first 2 d. When incubated for 3 d, the total diquat accumulation of *OsLAT1-OE* was approximately twice of that in wild type and *oslat1* mutants (Fig. 4B), explaining its increased dihydropyridine herbicide sensitivity.

**Fig. 4.**
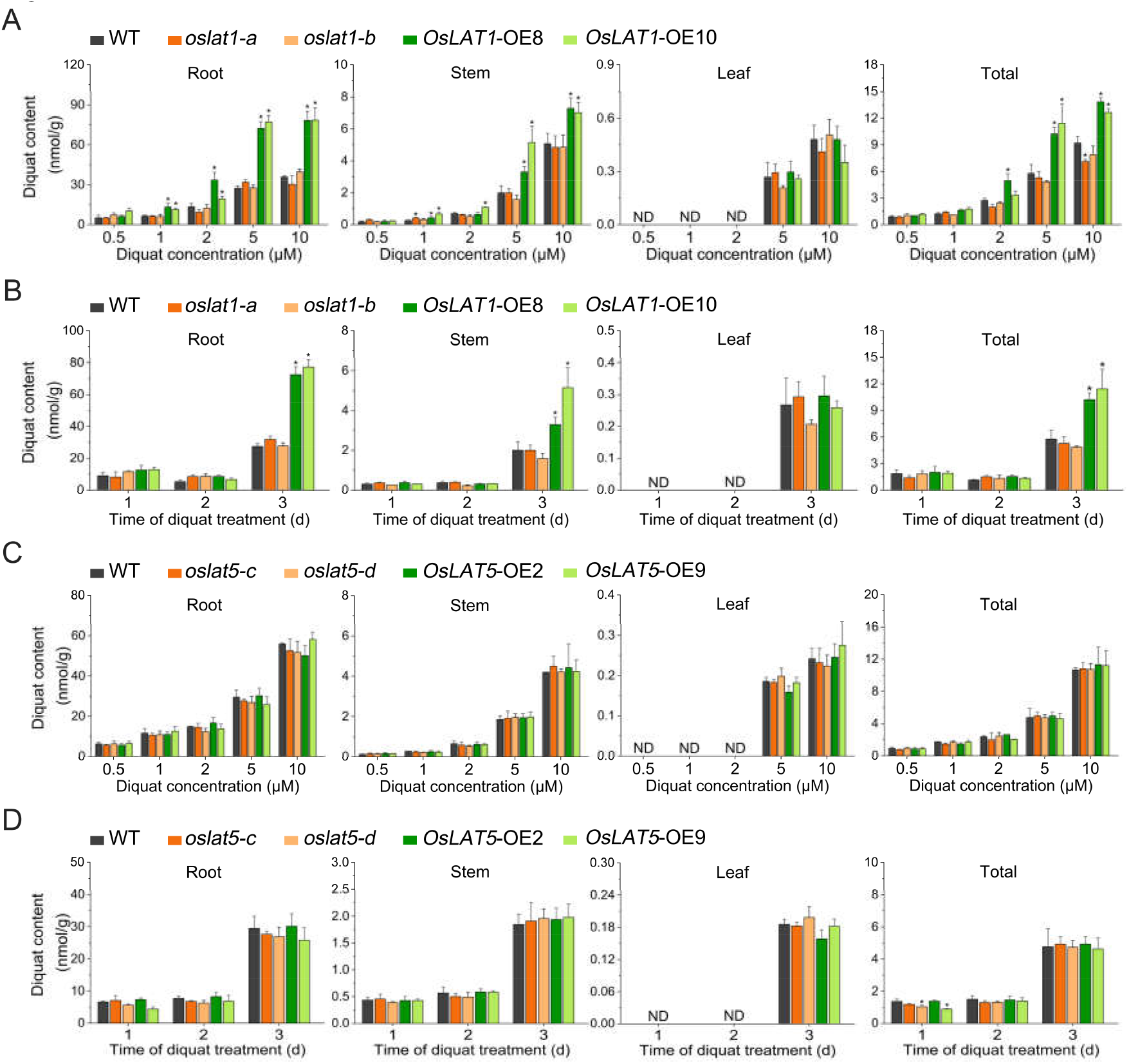
Measurement of diquat uptake in wild type, mutantation and overexpressing rice. (A and C) Dose-dependent diquat uptake. The seedlings of different lines of *OsLAT1* (A) or *OsLAT5* (C) (3 weeks old) were incubated with various concentration diquat for 3 days. (B and D) Time-dependent diquat uptake. The seedlings used were as those in (A) or (C), and were incubated with 5 μM diquat for the time period. Data are mean ± SD. n = 3. Asterisks indicate significant differences (one-way ANOVA: \**P* < 0.05).

On the other hand, there was no significant difference with respect to the diquat content among the *OsLAT5* rice genotype. Similarly, there was no difference in the diquat contents in different parts of rice plants (Fig. 4C and D). The uptake assays of another member of the dihydropyridine herbicide family, paraquat, were conducted as described for diquat. Paraquat uptake was similar to that of diquat (Fig. S7). This may have been linked to subcellular location of *OsLAT5*, meaning that it may have been responsible for transporting diquat into chloroplasts rather than working directly to achieve uptake. Brefeldin A (BFA) blocks the proteins intracellular trafficking to the Golgi apparatus by forming BFA compartmets. Therefore, in order to examine this possibility, we compared WT and *oslat5-c* with and without brefeldin A (BFA) in the MS medium (Fig. S8). The addition of BFA would have no effect upon the rice growth. However, it slightly improved resistance to dihydropyridine herbicide of *oslat5-c* mutant. These data supported the idea that *OsLAT5* may function in intracellular delivery of dihydropyridine herbicides, while *OsLAT1* is required for dihydropyridine herbicide uptake and transport from root to shoot.

### Application value of *OsLAT5* in innovating dihydropyridine herbicide-resistant rice

Based on the above results, *OsLAT5* played an important role in enhancing resistance to dihydropyridine herbicides. To explore the value of dihydropyridine herbicide-resistant rice with the *OsLAT5* mutation in the field, we generated the GY-*oslat5* mutant edited by CRISPR/Cas9 in the background Guiyu NO.11-a good-quality and high-yield cultivar (Fig. S9A). First, we investigated the function in dihydropyridine herbicide resistance by the seed germination and spray experiment. Under our assay condition, the *GY-oslat5* mutant remained tolerant to paraquat and diquat during germination and vegetative growth stages (Fig. S9B and C).

For the crop–weed competition assay, Guiyu NO.11 and the *GY-oslat5* mutant along with rice field weeds (e.g., *Echinochloa colona* L., *Cyperus rotundus* L., *Eclipta prostrata* L., and *Commelina communis* L.) were sown in a greenhouse, and the dihydropyridine herbicide was subsequently applied on one occasion. When herbicide was applied by either foliar spraying or irrigation, the GY11-*oslat5* mutant exhibited strong growth and was not affected under the dihydropyridine herbicide treatment. In contrast, Guiyu NO.11 wild-type plants were etiolating and appeared to have inhibited growth. It was also observed that dihydropyridine herbicide application resulted in mortality of *Cyperaceae (Cyperus rotundus* L.), broadleaf weeds (*Eclipta prostrata* L. and *Commelina communis* L.), and weak *Gramineae* individuals (*Echinochloa colona* L.) (Fig. 5). These results showed that the GY11-*oslat5* mutant exhibited a positive response to dihydropyridine herbicide application with foliar spraying and irrigation, and was effective in weed control to eliminate mainly weed species in rice cultivation.

**Fig. 5.**
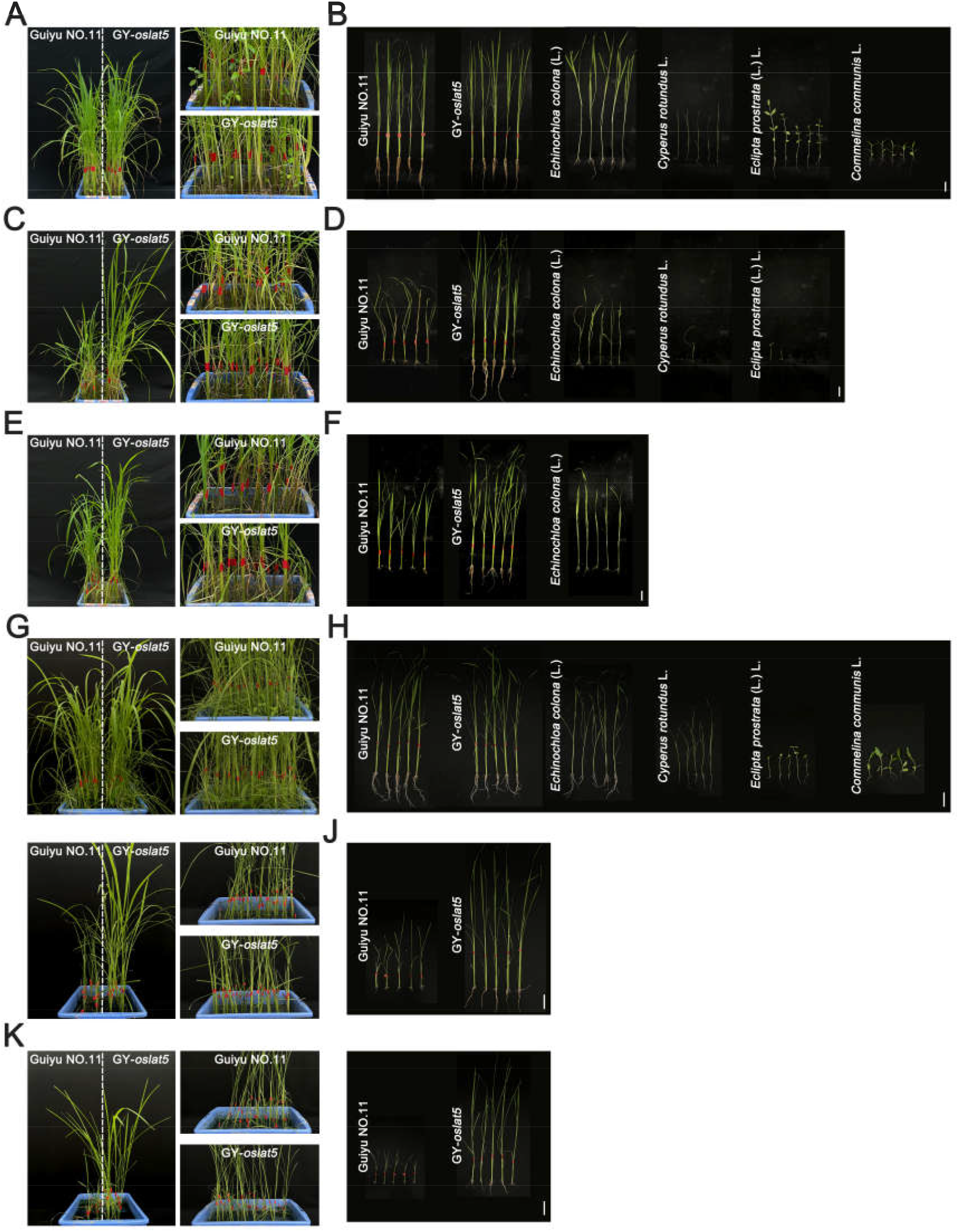
Evaluation of dihydropyridine herbicide on Guiyu NO.11, *GY-oslat5* mutant and weed by different application methods. (A-F) Effects of foliar spraying without herbicide (A-B) or with 5 mg/L paraquat (C-D) and 15 mg/L diquat (E-F) on rice and weeds grown in the soil for 12 days. The plants were taken pictures on the 28th day of dihydropyridine herbicide application. Bar = 5 cm. (G-L) Effects of irrigation with water (G-H), 2 mg/L paraquat (I-J) and 5 mg/L diquat (K-L) on rice and weeds grown in the soil for 5 days. Bar = 5 cm.

### Agronomic traits of dihydropyridine herbicide-resistant rice with *OsLAT5* mutation

The overall productive fitness and yield of wild-type Guiyu NO.11 and the *GY-oslat5* mutant were investigated under normal field conditions. The plant height of GY-*oslat5* decreased by ~8% compared to the wild type (Fig. 6A and C). Among the various studied yield parameters, panicle length and seed setting rate of GY-*oslat5* were comparable to those in the wild type (Fig. 6D and E), except that 1000-grain weight was reduced in GY-*oslat5* (Fig. 6F). However, a more detailed analysis on rice grains indicated no significant differences in length, width, or thickness between GY-*oslat5* and wild type (Fig. 6B), which may have been because of the grain characteristics and quality.

**Fig. 6.**
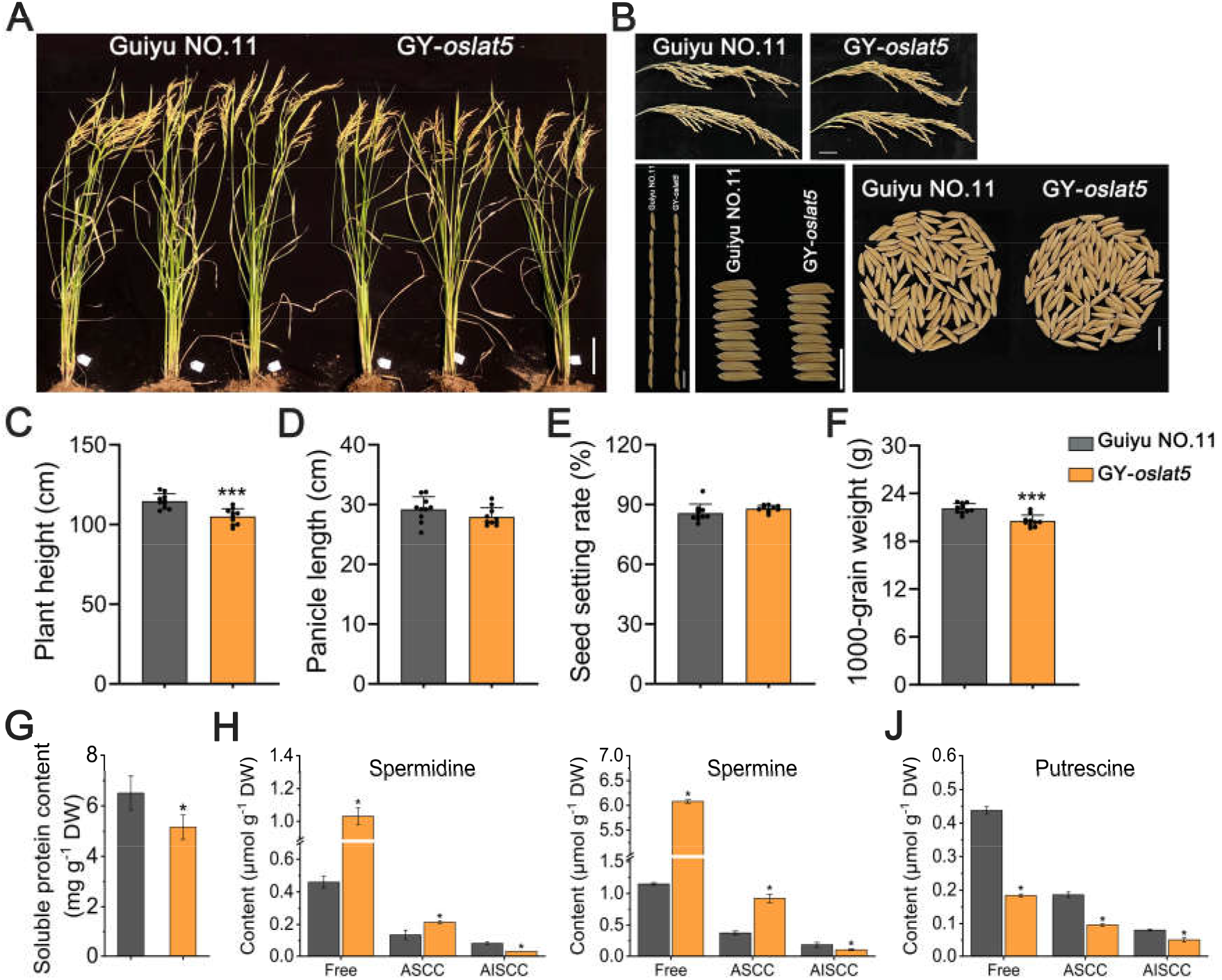
Agronomic traits analysis of *GY-oslat5* mutant compared with Guiyu NO.11. (A) The phenotypic difference between wild type and mutation of *OsLAT5* in normal field. Bar = 10 cm. (B) Phenotype of panicle and seed at maturation. Bar = 2 cm (panicle) or 1 cm (seed). (C-F) Comparison of plant height (C), panicle length (D), seed setting rate (E) and 1000-grain weight (F) between wild type and *GY-oslat5*. Data are mean ± SD. n = 10. Asterisks indicate significant differences (one-way ANOVA: \*\*\**P* < 0.001). (G) The content of soluble protein in the grain. (H-J) The content of spermidine (H), spermine (I) and putrescine (J) in the grain. Free, free polyamine; ASCC, acid soluble covalently conjugated polyamine; AISCC, acid insoluble covalently conjugated polyamine. Data are mean ± SD. n = 3. Asterisks indicate significant differences (one-way ANOVA: \**P* < 0.05).

Further analysis showed that the grain of GY-*oslat5* was higher in free or ASCC-spermidine (acid soluble covalently conjugated polyamine) and spermine, and lower in soluble protein, AISCC-spermidine (acid insoluble covalently conjugated polyamine) and spermine and all forms putrescine (Fig. 6G-6J). Thus, the knockdown of *OsLAT5* had a slightly reduction in plant length and 1000-grain weight, probably associated with the content of soluble protein and PAs in grain.

### 3D modelling reveals structural interaction of *OsLAT5* and dihydropyridine herbicide

According to the above-mentioned results, disruption of *OsLAT5* had negative impacts on rice growth, yields and changes of grain quality, it is of significant to validating the binding sites of *OsLAT5* and dihydropyridine herbicide instead of PA, which is necessary for rice yield and quality

Structural modelling predicted that OsLAT5 interacted with paraquat in the area including the amino acid residues Pro-44, Phe-45, and Phe-180, which provided hydrophobic interactions with paraquat (Fig. 7A). The binding model between OsLAT5 and diquat is shown in Fig. 7B. Diquat was located at a cavity formed by residues Phe-37, Ser-41, Pro-44, Phe-45, Phe-180, Tyr-252, Ser-298, and Asn-302. Among them, Ser-41, Ser-298, and Asn-302 formed hydrogen bond interactions to diquat, and the other residues provided hydrophobic interactions with diquat. We also found that Ser-50, Val-51, and Phe-451 may engage in OsLAT5 interactions (data not shown).

**Fig. 7.**
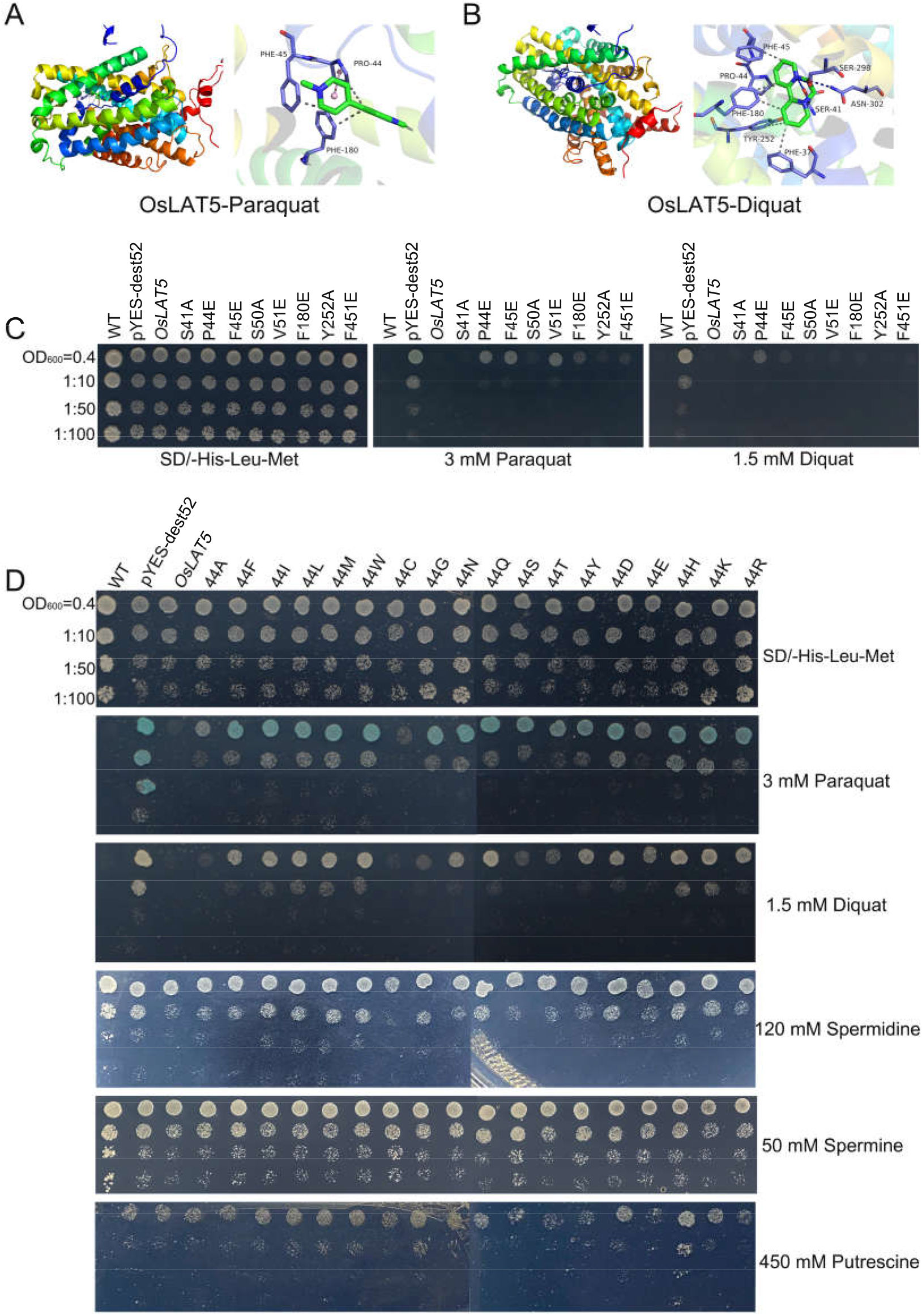
Structural and functional investigation of the dihydropyridine herbicide-binding residues of OsLAT5. (A-B) Spatial structure (left) of contact interface between OsLAT5 and paraquat (A) or diquat (B) and 2D-diagram (right) of intermolecular interactions. (C) Yeast cells were transformed with *OsLAT5* wild type or *OsLAT5* mutation. The mutations caused S41A, P44E, F45E, S50A, V51E, F180E, Y252A and F451E amino acid substitutions. The cells were grown on medium with 3 mM paraquat and 1.5 mM diquat or not for 5 days before pictures were taken. (D) Amino acid 44 of the OsLAT5 protein was mutated to various amino acid. The yeast transformants were grown on medium with paraquat (3 mM), diquat (1.5 mM), spermidine (120 mM), spermine (50 mM) and putrescine (450 mM) for 5 days before pictures were taken.

To identify the binding site of OsLAT5 and dihydropyridine herbicide, we mutated the amino acids involved in Ser-41, Pro-44, Phe-45, Phe-180, and Tyr-252 located in membrane-spanning regions, as well as Ser-50, Val-51, and Phe-451, to another different class of amino acid such as Ala or Glu. We then heterologously expressed the pYES-dest52 vector harboring the mutant gene in Δ*agp2*. It was cleared that the substituting Pro-44, Phe-45 and Val-51 with Glu abrogates the OsLAT5 activity in uptake paraquat, as the yeast strain expression them could survive on the drug-containing medium. Besides, only the strain carrying OsLAT5 with Pro-44-Glu mutant displayed tolerance to diquat. The implication of these results was that Pro-44 played an important role in the interaction between OsLAT5 and dihydropyridine herbicide.

Therefore, we continued to mutate the Pro-44 to 18 amino acids, then heterologously expressed the resulting proteins in yeast. Mutating Pro44 to Ala (44A), Cys (44C) and Glu (44E) did not change markedly the dihydropyridine herbicide transport activity of OsLAT5, other mutations including Gly (44G), Ser (44S) and Thr (44T) also could barely change on the medium with diquat. The rest may be the special sites for cultivating resistant rice with low dihydropyridine herbicide transport ability. As for the sensitivity to polyamine, only when Pro-44 was substituted with Phe, Tyr and Arg did the yeast remain sensitive to spermidine and putrescine (Fig. 7D). Together, Pro44 was essential binding sites of OsLAT5 for dihydropyridine herbicide transport, and the P44F, P44Y and P44R variant showed an enhancement in dihydropyridine herbicide resistance but no disruption of spermidine and putrescine transport competence, providing candidate sites for precise editing in breeding a dihydropyridine herbicide-resistant rice cultivar without impairing rice yield.

## Discussion

Most pesticides in contact with plants undergo the uptake and transport process. Uptake refers to the process of pesticides entering the plant, which can be achieved through the roots or leaves; whereas, pesticide transport in plants is generally divided into intracellular and inter-organ transport^45–48^. Recently, *OsATL15* was first identified to facilitate thiamethoxam uptake and systemic distribution in rice, which further confirmed that plant transporters may participate in pesticide entry into plants^49^. To take effect, herbicides must enter the plant, be translocated, and reach their target site^3^, and this process involves transporters. Many researchers have conducted studies in this area; for example, it has been shown that glyphosate could be exported to the extracellular space by ABC transporter (ABCC8) in *Echinochloa colona*^50^, and that paraquat was transported into cells by the PA transport system^24^. Dihydropyridine herbicide has been used worldwide for 50 y; however, it damages crop growth because of its non-selectivity, leading to limited applicability in orchards, plantation crops, and conservation tillage systems^51^. The concept of improving crops with herbicide tolerance by impairing transport activity via CRISPR/Cas9 technology is promising. Herbicide-resistant rice constructed by such means differs from the herbicide-tolerant genetically modified crops with low research and development efficiency and which are prone to gene escape^41^, and simplifies rice cultivation when combined with dihydropyridine herbicide application. Critical to the success of this approach is understanding the interaction underlying the herbicide and its transporters. In this study, we not only discovered that *OsLAT1* and *OsLAT5* are dihydropyridine herbicide transporters, but also dissected their functional differentiation in terms of transport functions in germination, seedling, and adult plant stages.

Although both *OsLAT1* and *OsLAT5* of four rice *LATs* could complement the Δ*agp2* phenotype of insensitivity to dihydropyridine herbicides (Fig. 1, S1, and S2), kinetic analysis showed that OsLAT5 had stronger affinity for dihydropyridine herbicides than OsLAT1 in yeast system. Furthermore, the expression pattern analysis of *OsLAT1* and *OsLAT5* along with semi-quantitative RT-PCR in rice tissues demonstrated that *OsLAT5* was remarkably higher than that of *OsLAT1* in various tissues. Furthermore, the signal of the GFP-tagged OsLAT1 was localized in the plasma membrane; whereas, OsLAT5 was in both the plasma membrane and Golgi apparatus (Fig. S3). The above evidence suggested that *OsLAT1* and *OsLAT5* fit the properties of dihydropyridine herbicide transporters, and that *OsLAT5* may play a greater role in transport of and tolerance to such herbicides.

At the plant level in rice, the root and shoot lengths of *OsLAT1*-OE rice were significantly inhibited by a low concentration of dihydropyridine herbicide (0.05 μM) during germination (Fig. 2C, 2E, S5C and S5E). This corresponded to the results of the dihydropyridine herbicide uptake experiment on rice seedlings. Excessive amounts of dihydropyridine herbicides were imported into *OsLAT1*-OE and then caused damage to plant growth, which contrasted to those of wild type and *OsLAT1* mutants. Moreover, *OsLAT1-OE* had higher paraquat and diquat contents in the roots and stems than that in the other two *OsLAT1* genotypes lines, indicating that overexpression of *OsLAT1* enhanced root uptake and the root-to-shoot transport ability of dihydropyridine herbicides in rice (Fig. 4A, 4B, S7A, and S7B). When commercial dihydropyridine herbicide concentrations were applied, differences between the wild type, *OsLAT1* mutant, and *OsLAT1*-OE were not reflected in rice growth and development, except that *OsLAT1*-OE conferred slightly more damage under the 200 mg/L diquat spraying treatment (Figs. 3A, 3B, S6A, and S6B). Overall, overexpression of *OsLAT1* enhanced the ability of dihydropyridine herbicide root uptake and transport to shoot; whereas, the mutation of *OsLAT1* did not exhibit a more resistant phenotype to dihydropyridine herbicide, which may have been attributed to genetic redundancy.

Regarding *OsLAT5*, the growth of the *oslat5* mutant at the rice germination stage exhibited no effect even under high dihydropyridine herbicide concentrations (Fig. 2D, 2F, S5D, S5F). After foliar spraying of dihydropyridine herbicide, the mutation of *OsLAT5* continued to play a role in resistance to these herbicides; surprisingly, the *oslat5* mutant gradually recovered to grow under all diquat concentration treatments (Figs. 3C, 3D, S6C, and S6D). However, we did not observe any significant difference in dihydropyridine herbicide uptake across *OsLAT5* rice genotypes (Fig. 4C, 4D, S7C, and S7D), implying that *OsLAT5* mainly functions in intracellular herbicide transport through a BFA-sensitive transport system (Fig. S8) as it is localized in the plasma membrane and Golgi apparatus (Fig. S3D). This view is supported by the observation made in a study on Golgi-localized transporter protein *PAR1*, in which paraquat tolerance of its mutant was attributed to a decrease in paraquat accumulation in the chloroplast instead of into plant^27^. Moreover, we applied *OsLAT5* with a superior performance in dihydropyridine herbicide resistance to create an herbicide-resistant rice variety. When sprayed or irrigated with dihydropyridine herbicide at the seedling stage, weak wild-type rice and weeds were observed, and only *GY-oslat5* mutants were healthy (Fig. 5). This variety exhibited a high application value in direct seeding rice fields. Furthermore, the applied dosage (maximum up to 15 mg/L) was lower than that used by farmers (approximately 50 mg/L), achieving the target of enhancing efficacy and reducing the pesticide application. Unfortunately, after diquat pelletization treatment, there was no significant control effect on the weeds compared with control treat, which was probably related to the passivation failure of diquat in soil (data not shown)

However, disruption of *OsLAT5* modestly affected rice yields and quality (Fig. 6). This result was expected since PA has been reported to be involved in plant development and biotic and abiotic stress responses^52^. Therefore, it is of significance to characterize the variants that exhibited substrate selectivity on dihydropyridine herbicide and PAs. Therefore, variants of P44F, P44Y and P44R of OsLAT5 was identified as important factor that could block transport dihydropyridine herbicide rather than spermidine and putrescine (Fig. 7D). And we observed that the exogenous application of spermidine and putrescine partly counteract dihydropyridine herbicide toxicity (Fig. S10), which means that the preservation of ability to transport polyamine helps to cultivate higher resistance rice. Besides, previous studies have shown that higher putrescine levels could promote spermidine and spermine synthesis and defend plants^52–54^. By site-specific editing this binding site of OsLAT5 and dihydropyridine herbicide, we can generate an iconic rice variety, which maintains resistance to such herbicides but not at the expense of poor growth or low grain yield.

In summary, OsLAT1 and OsLAT5 can act as dihydropyridine herbicide transport proteins; however, their functions are differentiated. OsLAT1, which is localized in the plasma membrane, is responsible for dihydropyridine herbicide root uptake and root-to-shoot transport; whereas, OsLAT5, localized in the plasma membrane and Golgi apparatus, is more responsible for intracellular transport of dihydropyridine herbicide to chloroplasts. Compared with the application values of genes *OsLAT1* and *OsLAT5* in generating dihydropyridine herbicide-resistant rice, *OsLAT5* contributed more and its variant carrying P44F, P44Y and P44R are expected to represent a promising candidate.

## Method

### Vector construction

The cloning vectors for *OsLAT1, OsLAT3, OsLAT5*, and *OsLAT7* were obtained from NARO DNA bank (https://www.dna.affrc.go.jp/). The PCR fragments amplified with gene-specific primers (Table S1) were sub-cloned into the expression vectors using the In-FfFusion kitmethod (Takara, Dalian, China), with the exception that the pYES-dest52 encoding target genes were constructed by LR recombination reaction (Invitrogen, Carlsbad, CA). All vectors were sequenced to confirm that inserts were correct.

### Yeast assay

The yeast mutant Δ*agp2* modified from wild-type BY4741 (*MATa his3Δ leu2Δ*, and *met15Δ ura3Δ*) was deficient in transporting PA. These two yeast strains were obtained from Dharmacon (*Δagp2* Catalog#: YSC6273-201931762, BY4741 Catalog#: YSC1048). The resulting pYES-dest52-*OsLAT1 OsLAT3, OsLAT5*, and *OsLAT7* constructs were transferred into the Δ*agp2* cells by using LiAc/SS-DNA/PEG method following the supplier’s protocol (Takara, Dalian, China), and the Δ*agp2* or BY4741 introduced into an empty pYES-dest52 vector and served as a control.

In the primary test, we screened the underlying dihydropyridine herbicide transporter genes. The yeast cells when harboring BY4741-empty vector, Δ*agp2*-empty vector, and Δ*agp2-candidate* genes were incubated on SD-Gal solid medium with histidine, leucine, methionine, and 2% glucose for 5 d at 30 °C. Pre-cultured cells were transferred to the SD-Gal (2% galactose) liquid medium and shaken at 250 rpm for 14–16 h. Cell suspensions were adjusted to an OD_600_ of 0.4 and then serially diluted with distilled water. Next, 3-μL aliquots were spotted to the solid SD-Gal containing either 0.75 mM paraquat or diquat, and plates were photographed after 5 d of incubation at 30°C. Dihydropyridine herbicide sensitive yeast was used for further experimental validation. We conducted the yeast growth assays on the solid medium following the same protocol; however, the difference was the SD-Gal with paraquat (0.5, 0.75, and 1.5 mM) or diquat (0.1, 0.5, and 0.75 mM). For the yeast growth in the liquid medium, the yeasts were shake-cultured in SD-Gal medium until an OD_600_ reading of 0.8 was obtained. SD-Gal liquid medium, containing paraquat and diquat (0.75 mM) or no herbicide, was then added to the cell cultures to obtain a 10x dilution. The cultures were shaken at 250 rpm and 30 °C. The OD_600_ of each culture was measured at 12, 24, 36, 48, 70, and 82 h using a plate reader, to plot the OD_600_-growth time function.

To characterize the transport capabilities of OsLAT1 or OsLAT5, we determined the kinetic constants *K*_m_ and *V*_max_ on dihydropyridine herbicide uptake. The transport assay was established as described^38^. The cultured cells were grown until an OD_600_ of 1.0 was measured in the SD-Gal liquid medium and were then harvested, washed three times with the uptake buffer (0.333 mM MES, pH 5.7, 2% galactose), and resuspended in the same buffer at a cell density of 5 × 10^7^ cells (100 μL^-1^). Next, 2 mL of each culture was transferred to 10 mL centrifuge tubes and uptake was initiated by the addition of various paraquat or diquat concentration (5, 10, 15, 20, 30, and 50 μM). Uptake was stopped 1 hour later, and tubes where then centrifuged at 3000 rpm for 5 min to discard the uptake buffer. After washing three times with ice cold uptake buffer to remove exogeneous drugs, cells were completely homogenized in methanol/water (1/1) via ultrasound and vortex. Finally, the mixture was evaporated with vacuum and resuspended in acetonitrile/water/formic acid (5/14/1) and analyzed by ultra-performance liquid chromatography-tandem mass spectrometric (UPLC-MS/MS). Michaelis–Menten parameters were obtained by a non-linear regression method using Origin 8.0 software (Microcal Software, Northampton, MA, USA).

### Plant materials and growth conditions

The study used two rice cultivars, Zhonghua11 and Guiyu NO.11, *A. thaliana* (variety Col-0), and agronomically important weeds species such as *Echinochloa colona* L., *Cyperus rotundus* L., *Eclipta prostrata* L., and *Commelina communis* L. *OsLAT1-a* and *OsLAT1-b* or *oslat5-c* and *oslat5-d* (in Zhongghua 11 background), edited via CRISPR/Cas9 system, were identified by sequencing PCR products amplified with gene-specific primers (Table S1). Different expression cassettes of *OsLAT1* and *OsLAT5* under the 35S promoter in pCAMBIA1300-35S were generated followed by transformation into Zhonghua11 with *Agrobacterium tumefaciens-edited* transformation. After overexpressed rice harboring the *OsLAT1* or *OsLAT5* gene were confirmed by 50 mg/L hygromycin tolerance and gene expression level analysis, two lines of *OsLAT1* or *OsLAT5 (OsLAT1-OE8* and *OsLAT1-*OE10 or *OsLAT5*-OE2 and *OsLAT5*-OE9) with relatively high expression levels were selected for further study.

Rice and weeds were grown in soil or modified Hoagland nutrient solution under greenhouse conditions with 14 h/10 h light/dark cycle, temperatures of 30 °C/28 °C in light/dark, and light intensity > 350 μmol m^-2^ s^-1^. Rice seedlings in MS medium (4.4 g/L M&S basal medium with vitamins, 30 g/L sucrose, and 3.5 g/L phytagel at a final pH of 5.8) were placed in the growth chamber (25 °C; 14 h/10 h light/dark cycle; 300 μmol m^-2^s^-1^ light intensity). *A. thaliana* were planted at 22 °C with 100 μmol m^-2^ s^-1^ under a 12:12 h light/dark photoperiod.

### Gene expression pattern analysis

Rice plants grown in soil under the greenhouse conditions were divided into root, stem, and leaf and collected 20 and 40 d after sowing. Total RNA from different part was isolated using a E.Z.N.A. Plant RNA Kit (Omega, Guangzhou, China) and reverse-transcribed into cDNA with a PrimeScript^™^ RT reagent Kit with gDNA Eraser (Takara, Dalian, China), and were then subjected to quantitative RT-PCR analysis using SYBR Premix Ex Taq II (Takara, Dalian, China). Primer sequences of genes and UBQ2 (NM_001402243.1) as the control were all enlisted in Table S1. The PCR-amplified products were detected on 1% agarose gel and photographed using the gal imaging system. Additionally, we investigated the microarray data using RiceXPro (https://ricexpro.dna.affrc.go.jp) and monitored the expression of OsLAT1 or OsLAT5 after 2, 4, and 9 h exposure to 100 μM dihydropyridine herbicide.

### Subcellular localization

The plasmid *OsLAT1-GFP* or *OsLAT5-GFP*, together with a plasma membrane marker (1008-mCherry) or Golgi marker (GmMan1-mCherry), were cotransformed into protoplasts, which were prepared from mature leaves of 5-week-old *Arabidopsis* plants^25^. Fluorescence was measured using excitation/emission wave lengths of 488/525 nm for GFP and 552/610 nm for mCherry with a Leica SP8 confocal laser scanning microscope.

### Dihydropyridine herbicide tolerance seed germination assay

Seeds were rinsed with 70% (v/v) ethanol for 3 min and then sodium hypochlorite (NaClO) 30% (w/v) for 30 min for surface disinfection, washed several times with sterile water, and then plated on MS medium. To investigate the effects of dihydropyridine herbicide on seed germination, 60 seeds were placed in MS medium supplemented with 0.5 μM paraquat or diquat. Seed germination was calculated when their radicle or germ length reached approximately 1 mm. To assess the effects on seedling morphology, seeds were germinated on MS medium containing paraquat or diquat (0.05 and 0.1 μM) for 2 weeks and seedling morphologies were recorded and photographed.

### Dihydropyridine herbicide spray assay

Under the spray method, 4-week-old plants were sprayed with different doses of commercial dihydropyridine herbicide (50, 100, or 200 mg/L), and the effects on plant morphology were observed. The chlorophyll level and maximum quantum yield of photosystem II (Fv/Fm) were measured simultaneously.

Total chlorophyll was extracted from randomly harvested leaves following the method described previously^25^. Fv/Fm was determined according to a previous study^55^ to confirm whether the photosynthetic performance suffered from paraquat or diquat exposures. After immersion in 0.1% agarose gel, the randomly collected leaves were placed in a dark room for at least 30 min, after which they were analyzed using a chlorophyll fluorometer (Mini PAM, Waltz, Effeltrich, Germany).

### Dihydropyridine herbicide uptake assay

The 3-week-old hydroponically grown rice were used for the root uptake assay following a method that was described previously, with modifications^27^. Briefly, seedlings were pre-treatment with uptake buffer consisting of 0.5 mM CaCl2 and 5 mM MES/Tris buffer (pH 5.8) for 60 min. In the time-course experiment, roots were incubated in 5 μM paraquat or diquat for 1–3 days. For the dose-dependent experiment, seedlings were exposed to paraquat or diquat concentration of 0.5–10 μM paraquat or diquat for 3 d. Roots, stems, and leaves were collected, weighed, and extracted with acetonitrile/water/formic acid (5:14:1) solution under ultrasonication and rotation. Next, stem and leaf samples were added to trichloromethane and vortexed vigorously for 3 min to remove pigments. Samples were then centrifuged at 3000 rpm for 10 min and the supernatant was allowed to vacuum dry before being re-dissolved in extract solution for UPLC-MS/MS analysis.

### Measurement of dihydropyridine herbicide level by UPLC-MS/MS

Chromatographic separation was carried out on an ACQUITY UPLC^®^ H-Class system (Waters) equipped with a CORTECS UPLC HILIC column (2.1 × 100 mm, 1.7 μm; Waters). The mobile phase was acetonitrile as mobile phase A and 10 mM ammonium acetate water with 1% formic acid as mobile phase B, which was performed at a flow of 0.4 mL/min. MS analysis was carried out on a Waters Xevo TQD triple-quadrupole mass spectrometer fitted with an ESI (electrospray ionization) source. The voltage parameters in the cone and capillary were set to 25 and 500 V, respectively, and the desolvation temperature was 500 °C. The following traces were monitored: paraquat, ESI+, m/z 185–158.1 (20 eV) for quantification, and m/z 185–169.7 (20 eV) for confirmation; diquat, ESI+, m/z 183.1–130.1 (30 eV) for quantification, and m/z 183.1–157.1 (25 eV) for confirmation.

### Application of dihydropyridine herbicide-resistant rice

A rice cultivar, Guiyu NO.11, as a high-yield and good-quality indica-type cultivar, was used for constructing dihydropyridine herbicide-resistant rice (GY-*oslat5*) using CRISPR/Cas9 technology. Following the confirmation of resistance to dihydropyridine herbicide, we tested crop–weed competition by planting rice lines (Guiyu NO.11 and GY-*oslat5*) along with weeds under the greenhouse conditions above-mentioned. All plants were exposed to dihydropyridine herbicide via one of three application methods: foliar spray, application in paddy field water layer, or seed-pelleting coating of herbicide.

Ten days after sowing in the soil, 5 mg/L paraquat or 15 mg/L diquat was sprayed on rice plants and weeds and symptoms were recorded daily.

Five days after sowing in the soil, the seedlings of rice and weeds were irrigated with water supplemented with 2 mg/L paraquat and 5 mg/L diquat, with 3 cm water was added to each treatment. The effects on plants were recorded.

The rice seeds, coating diquat in herbicide–seed ratios of 1:1000, 1:1500, and 1:2000, along with weeds were performed in the upper soil layer. The sowing density (50 seeds m^-2^) was chosen according to the rice direct seeding method. When rice shoots were 10 mm long, they were waterlogged to ensure a stable 10-mm-thick water layer above the soil surface.

### Field tests

To evaluate plant growth and rice yield in a natural environment, GY-*oslat5* and WT were grown in paddy rice fields in Guangzhou from February–June, 2022. The field was divided into six plots (i.e., three random plots per genotype), with each plot containing 3 rows × 20 plants per row. The growth parameters such as plant height, number of panicles per plant, panicle length, and yield were recorded at critical rice growing stages. For precision in the field test, rice plants growing on the edge were excluded to remove margin effects.

### Measurement of polyamine and soluble protein level in rice seed

Polyamines can be divided into free PAs, acid soluble covalently conjugated PAs (ASCC PAs) and acid insoluble covalently conjugated PAs (AISCC PAs)^56^. The extraction of all forms of polyamine was performed as described with modifications^57^. 0.2 g powder from seed sample were extracted with 2 mL 5% (v/v) perchloric acid, and placed in an ice bath for 1 h. Centrifugation of 15000×g for 20 min at 4°C. The supernatants were for the determination of free PAs and soluble conjugated PAs, and the sediments were for determination of insoluble bound PAs. For free PAs, collected 0.5 mL supernatants and added 1 mL NaOH (2 mol/L) to rotate, then mixed with 10 μL benzoyl chloride (CAS Number 98-88-4) and cultured at 30°C for 20 minutes for derivatizations. Adding 3 mL saturated sodium chloride and 3 ml of ether and forced to mix, then centrifugation of 20000×g for 10 min. Taking 0.5 mL of the supernatant and evaporating the organic solvent phase in vacuum, finally resuspended the residue in 0.5 mL of methanol for UPLC-MS/MS analysis. For ASCC PAs, collected 2 mL supernatants and added 2 mL HCl (12 mol/L) at 110°C for acidolysis. After 18 h, it was transferred to mortar and placed at 70°C and make sure the hydrochloric acid evaporated. Resuspending with 1 mL of 5% (v/v) perchloric acid and next for polyamines derivatizations as described above. For AISCC PAs, the sediments were added with 3 mL NaOH (1 mol/L) and 3 mL HCl (12 mol/L) for acidolysis. The next steps were as in ASCC PAs described. The benzylated polyamines were determined by UPLC-MS/MS. Chromatographic separation was carried out with an ACQUITY UPLC BEH C18 column (2.1 × 100 mm, 1.7 μm; Waters). The mobile phase 0.1% formic acid aqueous solution and acetonitrile were determined. Mass spectrometric parameters were the same as above of the measurement of dihydropyridine herbicide. The following traces were monitored: spermidine, ESI+, m/z 458.2-162.1 (25 eV) for quantification, m/z 458.2-105.2 (30 eV) for confirmation; spermine, ESI+, m/z 619.3-497.4 (20 eV) for quantification, m/z 619.3-105.2 (35 eV) for confirmation; putrescine, ESI+, m/z 297.2-105.2 (25 eV) for quantification, m/z 297.2-77 (40 eV) for confirmation. The soluble protein content in rice seed was measured by normal near-infrared method.

### Molecular docking of OsLAT5 with dihydropyridine herbicide

The spatial structure of OsLAT5 was predicted based on the web service “trRosetta”^58–60^. Paraquat and diquat structures were downloaded from PubChem (https://pubchem.ncbi.nlm.nih.gov/).

Molecular docking of paraquat and diquat molecules into the active site of OsLAT5 was performed via DOCK 6.9 software. The binding pocket of OsLAT5 were set as x = −6.6, 16.5, y = −4.3, 15.1, and z = −34.7, −16.1. After identification of docking pocket coordinates, docking was performed in the LeDock, followed by the picking of high-scoring docking from the output results for further analysis and preparation of the three-dimensional graphics using PyMOL1.7.

For studying the potential binding site, the point mutations in the pYES-dest52 vector were made using a Fast Site-Directed Mutagenesis kit (Tiangen, Beijing, China). The yeast growth on the solid medium assay referred to the above method. Besides dihydropyridine herbicide, we tested the tolerance of yeast cells to PAs including spermidine (75 mM), spermine (50 mM) and putrescine (400 mM).

## Supporting information

Supplemental informations

## Acknowledgments

We thank X.-Y He, Z.-H Lu, X.-S. Tian at Guangdong Academy of Agricultural Sciences for assistance with the assessment of agronomic traits and herbicide resistance in field. This work was funded by the National Natural Science Foundation of China (Grant No. 32072451) and Guangdong Modern Agricultural Industry Generic Key Technology Research and Development Innovation Team Project (Grant No. 2020KJ133), Generic Technique Innovation Team Construction of Modern Agriculture of Guangdong Province (2022KJ130), and the Project of Science and Technology in Guangdong Province (Grant No. 2018A030313188).

## Competing interests

For the application of *OsLAT5* and *OsLAT1* genes in rice breeding, Chinese patent licensing (No. ZL202010522403.8) were issued to the inventors H.-H. Xu., F. Lin and R.-H. Chen.

## Author contributions

The research was conceived and supervised by F. L. and H.-H. X.; R.-H. C., D.Z., C.-Z. L., D.-Y. Y, and S.-W. W. performed the experiments. R.-H. C. analyzed the data and drafted the manuscript. D.-Z., and C.-Z. L. prepared the figures. D.-Y. Y and S.-W. W. assisted in yeast screening, discussions during analysis, and reviewing the manuscript.

## Additional information

### Supplementary figure legends

**Fig. S1 The analysis of LAT family proteins in *A. thaliana* and *O. sativa*.**

(A) Phylogenetic trees depicting LAT family members in *A. thaliana* and *O. sativa*. (B) Growth of yeast mutant Δ*agp2* transformants carrying four *LATs* (1, 3, 5 and 7) genes. BY4741-empty vector (WT), Δ*agp2*-empty vector (pYES-dest52) and Δ*agp2-OsLATs* were incubated on SD/-His-Leu-Met (synthetic dropout medium only with histidine, leucine, methionine) solid medium supplemented with 2% galactose as well as 0.75 mM paraquat or diquat. The density of exponentially growing cell cultures was normalized to an OD_600_ of 0.4. Cell suspensions were serially diluted as indicated and 3 μL of each were spotted on the medium. Plates were photographed after 5 days of incubation at 30°C.

**Fig. S2 Dose-dependent uptake of dihydropyridine herbicide by *OsLAT1* and *OsLAT5*.**

Yeast cells (5×10^7^ cells/100 μL) including WT, pYES-dest52, *OsLAT1* and *OsLAT5* were incubated for 60 min in 0.333 mM MES (pH 5.7) and 2% galactose uptake buffer supplemented with 10, 20, 30, 40 and 50 μM paraquat (A) or diquat (B) to determine intracellular amount of paraquat or diquat. Data are mean ± SD. n = 3. Different letters indicate significant differences (*P* < 0.05; Duncan’s multiple range tests).

**Fig. S3 Expression pattern and subcellular localization of *OsLAT1* and *OsLAT5*.**

(A) Transcriptional analysis of *OsLAT1* and *OsLAT5* in root, stem and leaf of rice. Rice plants were grown in soil under the greenhouse conditions for the RNA isolation of different tissues. RT-PCR analysis were performed with gene-specific primers. Rice actin UBQ2 was used as internal control.

(B) Expression profiles of *OsLAT1* and *OsLAT5* from the public RiceXPro microarray data (https://ricexpro.dna.affrc.go.jp).

(C) Responses of *OsLAT1* or *OsLAT5* to dihydropyridine herbicide treatment. Four-week-old rice seedling were treated with 100 μM paraquat or diquat and then collected except root in the indicated time.

(D) Subcellular localization of OsLAT1 and OsLAT5 in *Arabidopsis* protoplasts.

**Fig. S4 Different lines of *OsLAT1* and *OsLAT5* in rice.**

(A-B) The *oslat1* and *oslat5* knockout mutants were generated by CRISPR-Cas9 technology, two different types of mutation were detected in genomic DNA of *oslat1-a* and *oslat1-b* (A) or *oslat5-c* and *oslat5-d* (B), respectively.

(C-D) Transcript levels of *OsLAT1* (C) and *OsLAT5* (D) under the control of 35S promoter were analyzed by qRT-PCR.

**Fig. S5 Rice seed germination and seedling growth on medium containing paraquat.**

(A) Seed germination curve. Seeds of wild type, mutants and over-expression line of *OsLAT1* or *OsLAT5* were germinated on MS medium added with 0 or 0.5 μM paraquat for 6 days and germination rate was recorded every day.

(B) The above seeds continued to grow followed by six days growing before pictures taken. Bar = 1 cm.

(C-D) Growth of germinated rice seedlings mediated by *OsLAT1* (C) or *OsLAT5* (D) with 0.05 and 0.1 μM paraquat on MS medium. Bar = 4 cm.

(E-F) The shoot and root length of rice seedlings shown in (C) and (D). Data are mean ± SD. n = 3. Asterisks indicate significant differences (one-way ANOVA:*\*P* < 0.05).

**Fig. S6 Effects of foliar spraying of paraquat on rice growth and development at tillering stage.**

(A) Field-grown rice of different lines of *OsLAT1* (4 weeks old) sprayed with 50, 100 and 200 mg/L paraquat followed by continue growth for different durations.

(B) Analysis of Fv/Fm and chlorophyll contents in the leaves from (A) treated with paraquat for the indicated durations. Data are mean ± SD. n =3. Asterisks indicate significant differences (one-way ANOVA: \**P* < 0.05).

(C) Field-grown rice of different lines of *OsLAT5* (4 weeks old) sprayed with 50, 100 and 200 mg/L paraquat followed by continue growth for different durations.

(D) Analysis of Fv/Fm and chlorophyll contents in the leaves from (C) treated with paraquat for the indicated durations. Data are mean ± SD. n = 3. Asterisks indicate significant differences (one-way ANOVA: \**P* < 0.05).

**Fig. S7 Measurement of paraquat uptake in wild type, mutant and overexpressing rice.**

(A and C) Dose-dependent paraquat uptake. The seedlings of different lines of *OsLAT1* (A) or *OsLAT5* (C) (3 weeks old) were incubated with various concentration paraquat for 3 days.

(B and D) Time-dependent paraquat uptake. The seedlings used were as those in (A) or (C), and were incubated with 5 μM paraquat for the time period.

Data are mean ± SD. n = 3. Asterisks indicate significant differences (one-way ANOVA: \**P* < 0.05).

**Fig. S8 Phenotypes of wild type and *oslat5-c* grown on MS plate (control) or MS plate containing 0.25 μM dihydropyridine herbicide and 0.25 μM dihydropyridine herbicide + 3 μM BFA.**

**Fig. S9 The *GY-oslat5* mutant phenotype.**

(A) The *oslat5* knockout mutants (background: Guiyu NO.11) were generated by CRISPR-Cas9 technology, the type of mutation was detected in genomic DNA of GY-*oslat5*.

(B) Effects of dihydropyridine herbicides in rice seed germination of *GY-oslat5*.

(C) Effects of foliar spraying of dihydropyridine herbicides on rice growth and development of GY-*oslat5*.

**Fig. S10 Effects of exogenous polyamines on dihydropyridine herbicide toxicity.**

(A) Phenotypes of Zhonghua11 grown on MS plate supplemented with 0.075 μM dihydropyridine herbicide and 1 mM polyamines.

(B) The shoot and root length of rice seedlings shown in (A). Data are mean ± SD. n = 3. Asterisks indicate significant differences (one-way ANOVA: \**P* < 0.05).

## Notes

### Competing Interest Statement

The authors have declared no competing interest.

## References

1. Kaspary, T. E., Roma-Burgos, N. & Merotto, A. Snorkeling Strategy: Tolerance to Flooding in Rice and Potential Application for Weed Management. Genes. 11, 975 (2020).

2. Deng, W. et al. Cyhalofop-butyl and Glyphosate Multiple-Herbicide Resistance Evolved in an *Eleusine indica* Population Collected in Chinese Direct-Seeding Rice. J. Agr. Food Chem. 68, 2623–2630 (2020).

3. Powles, S. B. & Yu, Q. Evolution in Action: Plants Resistant to Herbicides. Annu. Rev. Plant Biol. 61, 317–347 (2010).

4. Brian, R. C., Homer, R. F. & Stubbs, J. A new herbicide 1: 1-ethylene-2: 2-dipyrilium dibromide. Nature. 181, 446–447 (1958).

5. Funderburk, H. H. & Bozarth, G. A. Review of the metabolism and decompostion of diquat and paraquat. J. Agr. Food Chem. 15, 563–567 (1967).

6. Brian, R. C. Darkness and the activity of diquat and paraquat on tomato, broad bean and sugar beet. Ann. Appl. Biol. 60, 77–85 (1967).

7. Soar, C. J., Karotam, J., Preston, C. & Powles, S. B. Reduced paraquat translocation in paraquat resistant *Arctotheca calendula* (L.) Levyns is a consequence of the primary resistance mechanism, not the cause. Pestic. Biochem. Phys. 76, 91–98 (2003).

8. Gondar, D., López, R., Antelo, J., Fiol, S. & Arce, F. Adsorption of paraquat on soil organic matter: Effect of exchangeable cations and dissolved organic carbon. J. Hazard. Mater. 235-236, 218–223 (2012).

9. Hayes, M., Pick, M. E. & Toms, B. A. Interactions between clay minerals and bipyridylium herbicides. Residue Reviews. 57, 1–25 (1975).

10. Sartori, F. & Vidrio, E. Environmental fate and ecotoxicology of paraquat: a California perspective. Toxicological & Environmental Chemistry. 100, 479–517 (2018).

11. An, J. et al. Transcriptome profiling to discover putative genes associated with paraquat resistance in goosegrass *(Eleusine indica* L.). PLoS One. 9, e99940 (2014).

12. Farrington, J. A., Ebert, M., Land, E. J. & Fletcher, K. Bipyridylium quaternary salts and related compounds. V. Pulse radiolysis studies of the reaction of paraquat radical with oxygen. Implications for the mode of action of bipyridyl herbicides. Biochimica et Biophysica Acta (BBA)-Bioenergetics. 314, 372–381 (1973).

13. Dinis-Oliveira, R. J. et al. Paraquat poisonings: mechanisms of lung toxicity, clinical features, and treatment. Crit. Rev. Toxicol. 38, 13–71 (2008).

14. Rose, M. S., Smith, L. L. & Wyatt, I. Evidence for energy-dependent accumulation of paraquat into rat lung. Nature. 252, 314–315 (1974).

15. Hart, J. J., Di Tomaso, J. M., Linscott, D. L. & Kochian, L. V. Characterization of the transport and cellular compartmentation of paraquat in roots of intact maize seedlings. Pestic. Biochem. Phys. 43, 212–222 (1992).

16. Chang, C. J. & Kao, C. H. Paraquat toxicity is reduced by polyamines in rice leaves. Plant Growth Regul. 22, 163–168 (1997).

17. Hart, J. J., DiTomaso, J. M., Linscott, D. L. & Kochian, L. V. Transport interactions between paraquat and polyamines in roots of intact maize seedlings. Plant Physiol. 99, 1400–1405 (1992).

18. Soar, C. J., Preston, C., Karotam, J. & Powles, S. B. Polyamines can inhibit paraquat toxicity and translocation in the broadleaf weed *Arctotheca calendula*. Pestic. Biochem. Phys. 80, 94–105 (2004).

19. Ye, B., Muller, H. H., Zhang, J. & Gressel, J. Constitutively Elevated Levels of Putrescine and Putrescine-Generating Enzymes Correlated with Oxidant Stress Resistance in *Conyza bonariensis* and Wheat. Plant Physiol. 115, 1443–1451 (1997).

20. Benavides, M. P., Gallego, S. M., Comba, M. E. & Tomaro, M. L. Relationship between polyamines and paraquat toxicity in sunflower leaf discs. Plant Growth Regul. 31, 215–224 (2000).

21. Ross, J. H. & Krieger, R. I. Structure-activity correlations of amines inhibiting active uptake of paraquat (methyl viologen) into rat lung slices. Toxicol. Appl. Pharm. 59, 238–249 (1981).

22. Gordonsmith, R. H., Brooke-Taylor, S., Smith, L. L. & Cohen, G. M. Structural requirements of compounds to inhibit pulmonary diamine accumulation. Biochem. Pharmacol. 32, 3701–3709 (1983).

23. Fujita, M. et al. Natural variation in a polyamine transporter determines paraquat tolerance in Arabidopsis. Proc. Natl. Acad. Sci. USA 109, 6343–6347 (2012).

24. Fujita, M. & Shinozaki, K. Identification of Polyamine Transporters in plants: Paraquat transport provides crucial clues. Plant Cell Physiol. 55, 855–861 (2014).

25. Li, J. et al. Paraquat Resistant1, a Golgi-localized putative transporter protein, is involved in intracellular transport of paraquat. Plant Physiol. 162, 470–483 (2013).

26. Dong, S. et al. A pqr2 mutant encodes a defective polyamine transporter and is negatively affected by ABA for paraquat resistance in *Arabidopsis thaliana*. J. Plant Res. 129, 899–907 (2016).

27. Xi, J., Xu, P. & Xiang, C. Loss of AtPDR11, a plasma membrane-localized ABC transporter, confers paraquat tolerance in *Arabidopsis thaliana*. The Plant Journal 69, 782–791 (2012).

28. Luo, Q., Wei, J., Dong, Z., Shen, X. & Chen, Y. Differences of endogenous polyamines and putative genes associated with paraquat resistance in goosegrass *(Eleusine indica L.)*. PLoS One. 14, e216513 (2019).

29. Wen, X. et al. MDR1 Transporter protects against paraquat-induced toxicity in human and mouse proximal tubule cells. Toxicol. Sci. 141, 475–483 (2014).

30. Xia, J. et al. A gain-of-function mutation of the MATE family transporter DTX6 confers paraquat resistance in Arabidopsis. Mol. Plant. 14, 2126–2133 (2021).

31. Lv, Z. et al. Changing Gly311 to an acidic amino acid in the MATE family protein DTX6 enhances *Arabidopsis* resistance to the dihydropyridine herbicides. Mol. Plant 14, 2115–2125 (2021).

32. Begam, R. A. & Good, A. G. The *Arabidopsis* paraquat resistant1 mutant accumulates leucine upon dark treatment. Botany 95, 751–761 (2017).

33. Okumoto, S. & Pilot, G. Amino acid export in plants: A missing link in nitrogen cycling. Mol. Plant 4, 453–463 (2011).

34. Wipf, D. et al. Conservation of amino acid transporters in fungi, plants and animals. Trends Biochem Sci. 27, 139–147 (2002).

35. Uchino, H. et al. Transport of amino acid-related compounds mediated by L-type amino acid transporter 1 (LAT1): insights into the mechanisms of substrate recognitionz. Mol. Pharmacol. 61, 729–737 (2002).

36. van Veen, S. et al. ATP13A2 deficiency disrupts lysosomal polyamine export. Nature. 578, 419–424 (2020).

37. Mulangi, V., Phuntumart, V., Aouida, M., Ramotar, D. & Morris, P. Functional analysis of *OsPUT1,* a rice polyamine uptake transporter. Planta. 235, 1–11 (2012).

38. Mulangi, V., Chibucos, M. C., Phuntumart, V. & Morris, P. F. Kinetic and phylogenetic analysis of plant polyamine uptake transporters. Planta. 236, 1261–1273 (2012).

39. Lyu, Y., Cao, L., Huang, W., Liu, J. & Lu, H. Disruption of three polyamine uptake transporter genes in rice by CRISPR/Cas9 gene editing confers tolerance to herbicide paraquat. aBIOTECH. 3, 140–145 (2022).

40. Gao, C. Genome engineering for crop improvement and future agriculture. Cell. 184, 1621–1635 (2021).

41. Jin, M., Chen, L., Deng, X. W. & Tang, X. Development of herbicide resistance genes and their application in rice. The Crop Journal. 10, 26–35 (2021).

42. Igarashi, K. & Kashiwagi, K. Characteristics of cellular polyamine transport in prokaryotes and eukaryotes. Plant Physiol. Bioch. 48, 506–512 (2010).

43. Bishop, T., Powles, S. B. & Cornic, G. Mechanism of Paraquat Resistance in Hordeum glaucum. 11.* Paraquat Uptake and Translocation. Funct. Plant Biol. 14, 539–547 (1987).

44. Lasat, M. M., Ditomaso, J. M., Hart, J. J. & Kochian, L. V. Evidence for vacuolar sequestration of paraquat in roots of a paraquat-resistant Hordeum glaucum biotype. Physiol. Plantarum. 99, 255–262 (2010).

45. Satchivi, N. M., Stoller, E. W., Wax, L. M. & Briskin, D. P. A nonlinear dynamic simulation model for xenobiotic transport and whole plant allocation following foliar application. II. Model validation. Pestic. Biochem. Phys. 68, 67–84 (2000).

46. Qiu, J. et al. In vivo tracing of organochloride and organophosphorus pesticides in different organs of hydroponically grown malabar spinach *(Basella alba* L.). J. Hazard. Mater. 316, 52–59 (2016).

47. Wu, H., Xu, H., Marivingt Mounir, C., Bonnemain, J. L. & Chollet, J. F. Vectorizing agrochemicals: enhancing bioavailability via carrier-mediated transport. Pest Manag. Sci. 75, 1507–1516 (2019).

48. Park, J., Lee, Y., Martinoia, E. & Geisler, M. Plant hormone transporters: what we know and what we would like to know. BMC Biol. 15, 93 (2017).

49. Xiao, Y. et al. An amino acid transporter-like protein (OsATL15) facilitates the systematic distribution of thiamethoxam in rice for controlling the brown planthopper. Plant Biotechnol. J. (2022). DOI: 10.1111/pbi.13869.

50. Pan, L. et al. An ABCC-type transporter endowing glyphosate resistance in plants. Proc. Natl. Acad. Sci. USA 118, e2100136118 (2021).

51. Bromilow, R. H. Paraquat and sustainable agriculture. Pest Manag. Sci. 60, 340–349 (2004).

52. Gerlin, L., Baroukh, C. & Genin, S. Polyamines: double agents in disease and plant immunity. Trends Plant Sci. 26, 1061–1071 (2021).

53. Fuell, C., Elliott, K. A., Hanfrey, C. C., Franceschetti, M. & Michael, A. J. Polyamine biosynthetic diversity in plants and algae. Plant Physiol. Bioch. 48, 513–520 (2010).

54. Galston, A. W. & Sawhney, R. K. Polyamines in plant physiology. Plant Physiol. 94, 406–410 (1990).

55. Fu, X. K., Liu, C., Li, X. L. & Wang, G. Determination of the maximum quantum yield of PS II photochemistry in higher plants. Bio-101. e1010168 (2018). DOI: 10.21769/BioProtoc.1010168.

56. Sen, K., Choudhuri, M. M. & Ghosh, B. Changes in polyamine contents during development and germination of rice seeds. Phytochemistry. 20, 631–633 (1981).

57. Kiriakos, K., Maria, D., Christakis, H., Kalliopi, A. & Roubelakis, A. A narrow-pore HPLC method for the identification and quantitation of free, conjugated, and bound polyamines. Anal. Biochem. 214, 484–489 (1993).

58. Du, Z. et al. The trRosetta server for fast and accurate protein structure prediction. Nat. Protoc. 16, 5634–5651 (2021).

59. Su, H. et al. Improved protein structure prediction using a new multi-scale network and homologous templates. Advanced Science. 8, 2102592 (2021).

60. Yang, J. et al. Improved protein structure prediction using predicted interresidue orientations. Proc. Natl. Acad. Sci. USA 117, 1496–1503 (2020).

