## Supplemental informations for "Disruption of *OsLAT5* is sufficient to endow rice tolerance to dihydropyridine herbicides at commercial application concentrations"

### Supplementary table

**Table S1 Primers sequences used in this study.**

| Name | Primer sequence | Vector |
|---|---|---|
| pENTR-*OsLAT1* | P1: **ACCAATTCAGTCGACTGGATCC**ATGGCGGACACCGGCGGAC | |
| | P2: **TACAAGAAAGCTGGGTCTAGATATC**CTACGGAACCAACGGCTCG | pENTR 2B Dual |
| pENTR-*OsLAT5* | P3: **ACCAATTCAGTCGACTGGATCC**ATGGAGGATTGTGTTGGTAT | |
| | P4: **TACAAGAAAGCTGGGTCTAGATATC**TCAGCACACAAGTGGGATT | |
| qRT-PCR-*OsLAT1* | P5: ATCGGCTTCTTGGTCCTCC | |
| | P6: CCACTTCATCCACCCTTGC | |
| qRT-PCR-*OsLAT5* | P7: CGCCTCCCGACCTTACAA | |
| | P8: GCACGAACCCAACCAGCA | |
| Actin (UBQ2) | P9: TGCTATGTACGTCGCCATCCAG | |
| | P10: AATGAGTAACCACGCTCCGTCA | |
| GFP-*OsLAT1* | P11: **GCTGTACAAGCGAGCTCAAGCTTCG**ATGGCGGACACCGGCGGAC | |
| | P12: **TGCGGACTCTAGATCAGGTGGATCC**CGGAACCAACGGCTCG | pYL322-d1-eGFPc |
| GFP-*OsLAT5* | P13: **GCTGTACAAGCGAGCTCAAGCTTCG**ATGGAGGATTGTGTTGGTAT | |
| | P14: **TGCGGACTCTAGATCAGGTGGATCC**GCACACAAGTGGGATT | |
| 35S-*OsLAT1* | P15: **ACCCGGGGATCCTCTAGAGTCGA**ATGGCGGACACCGGCGGAC | |
| | P16: **ATGATACGAACGAAAGCTCTGCA**CTACGGAACCAACGGCTCG | pCAMB1300-35s |
| 35S-*OsLAT5* | P17: **ACCCGGGGATCCTCTAGAGTCGA**ATGGAGGATTGTGTTGGTAT | |
| | P18: **ATGATACGAACGAAAGCTCTGCA**TCAGCACACAAGTGGGATT | |
| Crispr-*OsLAT1* | P19: GTAGCGATGCGACGAGAGAG | |
| | P20: AAATGTGCTGCGCGCTTTAT | |
| Crispr-*OsLAT5* | P21: GTGTTGCATGGTGAAACAAGGA | |
| | P22: CAAGGGCTGAAGAGACCCAG | |
| S41A-*OsLAT5* | P23: TTTGGGATTGAGGATAGTGTC | |
| | P24: CGGACCCCCAGCAACTTCATA | |
| P44E-*OsLAT5* | P25: GAAGTTTCTGGGGGTGAGTTTGGGA | |
| | P26: ATAGAATATGAGGAAAATGAGTGGG | |
| F45E-*OsLAT5* | P27: TCTGGGGGTCCGGAGGGGATTGA | |
| | P28: AACTTCATAGAATATGAGGAAAATG | |
| S50A-*OsLAT5* | P29: GGATTGAGGATGCTGTCAAGGCTGC | |
| | P30: GCAGCCTTGACAGCATCCTCAATCC | |
| V51E-*OsLAT5* | P31: GAGGATAGTGAGAAGGCTGCTGGC | |
| | P32: GCCAGCAGCCTTCTCACTATCCTC | |
| F180E-*OsLAT5* | P33: GGATTAATAGCTATTCCCCGAA | |
| | P34: CATAACAAACTCCGGGAGTAGAGAG | |
| Y252A-*OsLAT5* | P35: TGGTGGGGGGAGCCCTCTACCCT | |
| | P36: CTAAAACTAGAGCATAAGAAA | |
| F451E-*OsLAT5* | P37: CGGTGGCTGAAGGAGTCCATAAGCGCAG | |
| | P38: CTGCGCTTATGGACTCCTTCAGCCACCG | |
| 44A-*OsLAT5* | P39: AAGTTTCTGGGGGTGCGTTTGGGATTGA | |
| | P40: CACCCCCAGAAACTTCATAGAATATGAG | pYES-dest52 |
| 44F-*OsLAT5* | P41: GAAGTTTCTGGGGGTTTCTTTGGGATTGAGGAT | |
| | P42: ATCCTCAATCCCAAAGAAACCCCCAGAAACTTC | |
| 44G-*OsLAT5* | P43: AAGTTTCTGGGGGTGGGTTTGGGATTGAGG | |
| | P44: CCTCAATCCCAAACCCACCCCCAGAAACTT | |
| 44I-*OsLAT5* | P45: GAAGTTTCTGGGGGTATATTTGGGATTGAGGAT | |
| | P46: ATCCTCAATCCCAAATATACCCCCAGAAACTTC | |
| 44L-*OsLAT5* | P47: AAGTTTCTGGGGGTCTGTTTGGGATTGAGGA | |
| | P48: AGACCCCCAGAAACTTCATAGAATATGAGG | |
| 44M-*OsLAT5* | P49: GAAGTTTCTGGGGGTATGTTTGGGATTGAGGA | |
| | P50: TCCTCAATCCCAAACATACCCCCAGAAACTTC | |
| 44W-*OsLAT5* | P51: GAAGTTTCTGGGGGTTGGTTTGGGATTGAGGA | |
| | P52: TCCTCAATCCCAAACCAACCCCCAGAAACTTC | |
| 44C-*OsLAT5* | P53: GAAGTTTCTGGGGGTTGCTTTGGGATTGAGGAT | |
| | P54: ATCCTCAATCCCAAAGCAACCCCCAGAAACTTC | |
| 44N-*OsLAT5* | P55: GAAGTTTCTGGGGGTAATTTTGGGATTGAGGAT | |
| | P56: ATCCTCAATCCCAAAATTACCCCCAGAAACTTC | |

**Countiue:**

| Name | Primer sequence | Vector |
| --- | --- | --- |
| 44Q-*OsLAT5* | P57: AGTTTCTGGGGGTCAGTTTGGGATTGAGG | |
| | P58: TGACCCCCAGAAACTTCATAGAATATGA | |
| 44S-*OsLAT5* | P59: AAGTTTCTGGGGGTTCGTTTGGGATTGAG | |
| | P60: AACCCCCAGAAACTTCATAGAATATGAG | |
| 44T-*OsLAT5* | P61: AAGTTTCTGGGGGTACGTTTGGGATTGAG | |
| | P62: TACCCCCAGAAACTTCATAGAATATGAGG | |
| 44Y-*OsLAT5* | P63: GAAGTTTCTGGGGGTTATTTTGGGATTGAGGAT | |
| | P64: ATCCTCAATCCCAAAATAACCCCCAGAAACTTC | |
| 44D-*OsLAT5* | P65: AAGTTTCTGGGGGTGATTTTGGGATTGAGGA | pYES-dest52 |
| | P66: TCCTCAATCCCAAAATCACCCCCAGAAACTT | |
| 44H-*OsLAT5* | P67: AGTTTCTGGGGGTCATTTTGGGATTGAGGA | |
| | P68: TCCTCAATCCCAAAATGACCCCCAGAAACT | |
| 44K-*OsLAT5* | P69: GAAGTTTCTGGGGGTAAGTTTGGGATTGAGGA | |
| | P70: TCCTCAATCCCAAACTTACCCCCAGAAACTTC | |
| 44R-*OsLAT5* | P71: GTTTCTGGGGGTCGGTTTGGGATTGAGGA | |
| | P72: CGACCCCCAGAAACTTCATAGAATATGAG | |

Figure.S1

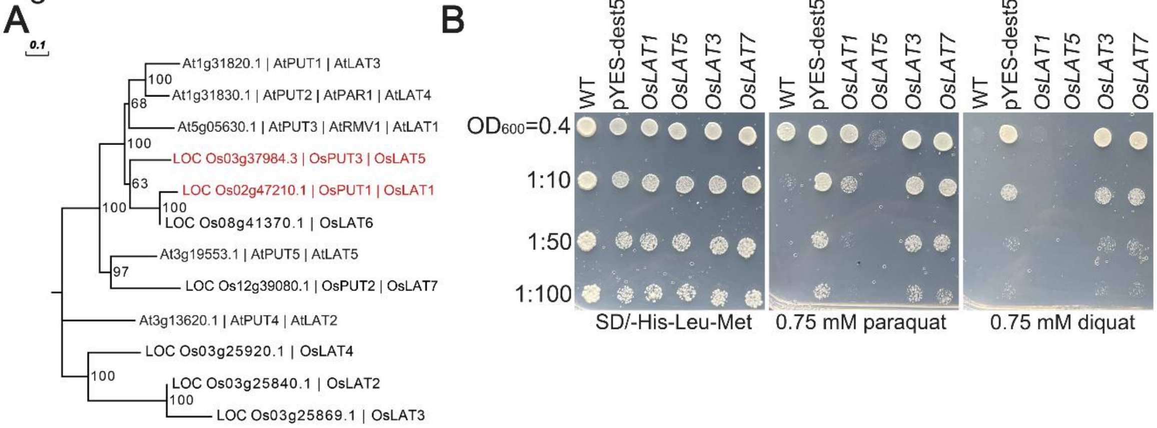


**Fig. S1 The analysis of LAT family proteins in *A. thaliana* and *O. sativa*.**

(A) Phylogenetic trees depicting LAT family members in *A. thaliana* and *O. sativa.*

(B) Growth of yeast mutant Δ*agp2* transformants carrying four *LATs* (1, 3, 5 and 7) genes. BY4741-empty vector (WT), Δ*agp2*-empty vector (pYES-dest52) and Δ*agp2*-*OsLATs* were incubated on SD/-His-Leu-Met (synthetic dropout medium only with histidine, leucine, methionine) solid medium supplemented with 2% galactose as well as 0.75 mM paraquat or diquat. The density of exponentially growing cell cultures was normalized to an $OD_{600}$ of 0.4. Cell suspensions were serially diluted as indicated and 3 μL of each were spotted on the medium. Plates were photographed after 5 days of incubation at 30°C.

Figure.S2

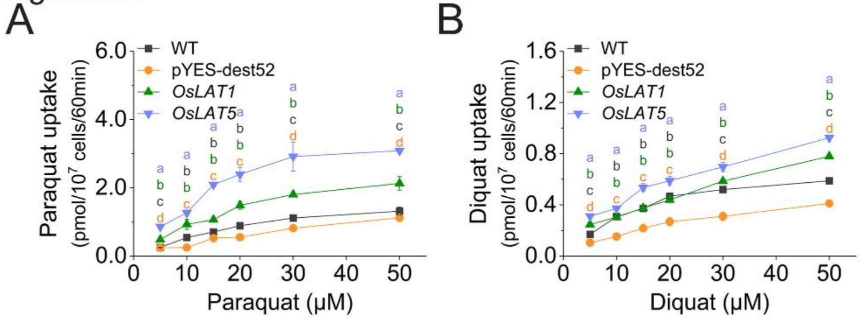


**Fig. S2 Dose-dependent uptake of dihydropyridine herbicide by *OsLAT1* and *OsLAT5*.** Yeast cells ($5\times10^7$ cells/100 μL) including WT, pYES-dest52, *OsLAT1* and *OsLAT5* were incubated for 60 min in 0.333 mM MES (pH 5.7) and 2% galactose uptake buffer supplemented with 10, 20, 30, 40 and 50 μM paraquat (A) or diquat (B) to determine intracellular amount of paraquat or diquat. Data are mean ± SD. n = 3. Different letters indicate significant differences ($P < 0.05$; Duncan's multiple range tests).

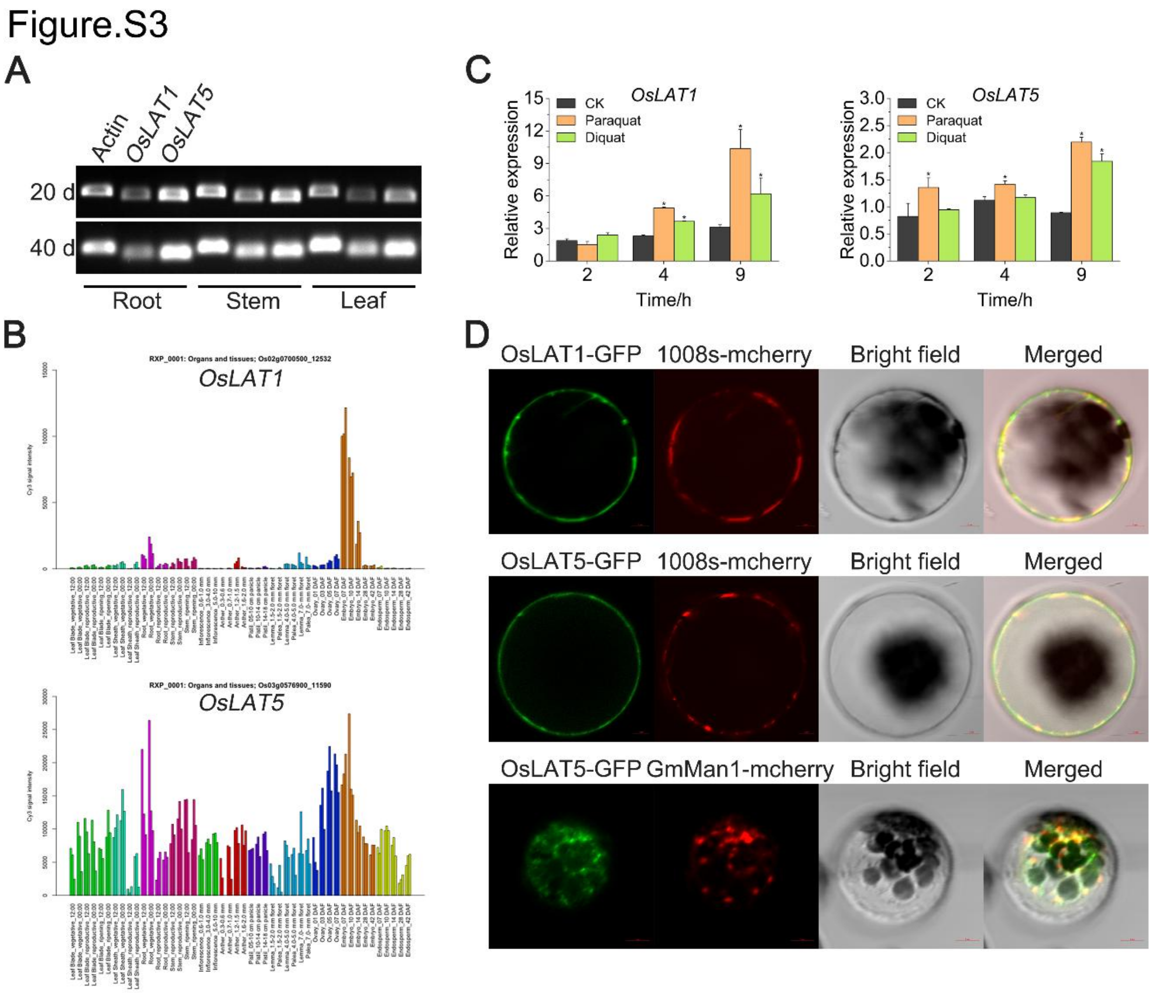


**Fig. S3 Expression pattern and subcellular localization of *OsLAT1* and *OsLAT5*.**

(A) Transcriptional analysis of *OsLAT1* and *OsLAT5* in root, stem and leaf of rice. Rice plants were grown in soil under the greenhouse conditions for the RNA isolation of different tissues. RT-PCR analysis were performed with gene-specific primers. Rice actin UBQ2 was used as internal control.

(B) Expression profiles of *OsLAT1* and *OsLAT5* from the public RiceXPro microarray data (https://ricexpro.dna.affrc.go.jp).

(C) Responses of *OsLAT1* or *OsLAT5* to dihydropyridine herbicide treatment. Four-week-old rice seedling were treated with 100 μM paraquat or diquat and then collected except root in the indicated time.

(D) Subcellular localization of OsLAT1 and OsLAT5 in *Arabidopsis* protoplasts.

Figure.S4

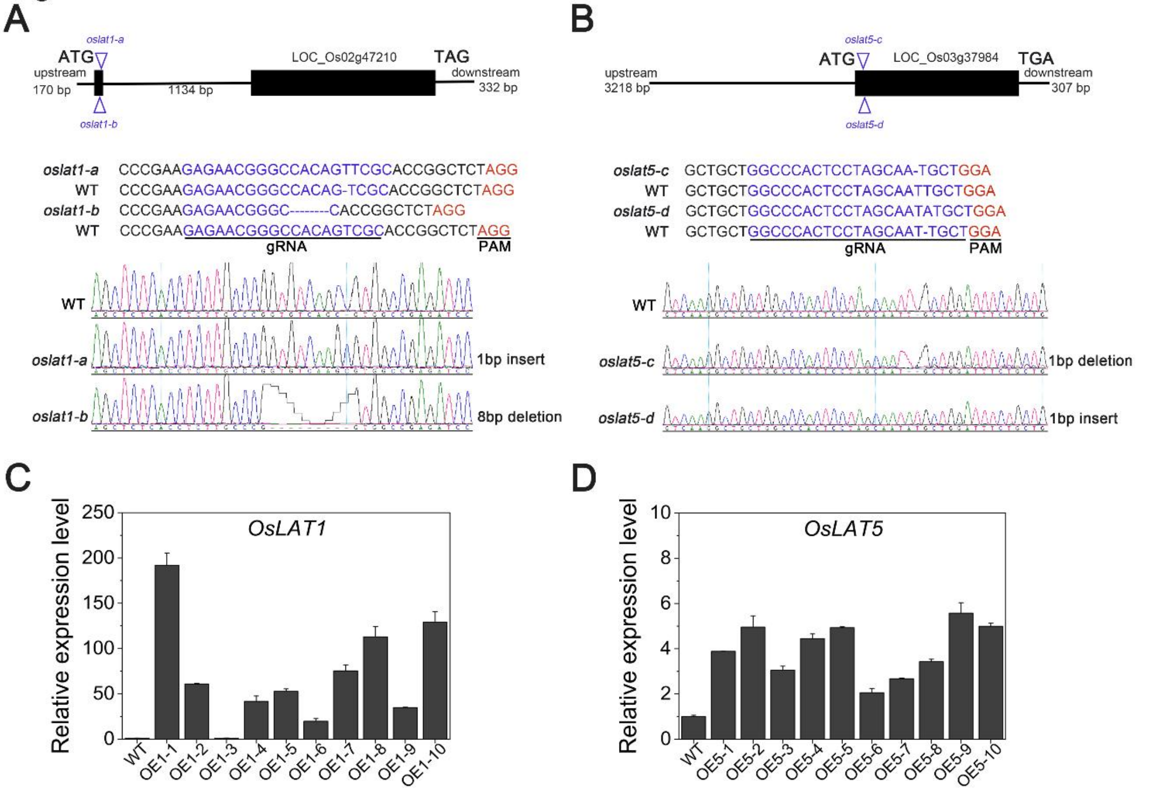


**Fig. S4 Different lines of *OsLAT1* and *OsLAT5* in rice.**

(A-B) The *oslat1* and *oslat5* knockout mutants were generated by CRISPR-Cas9 technology, two different types of mutation were detected in genomic DNA of *oslat1-a* and *oslat1-b* (A) or *oslat5-c* and *oslat5-d* (B), respectively.

(C-D) Transcript levels of *OsLAT1* (C) and *OsLAT5* (D) under the control of 35S promoter were analyzed by qRT-PCR.

Figure.S5

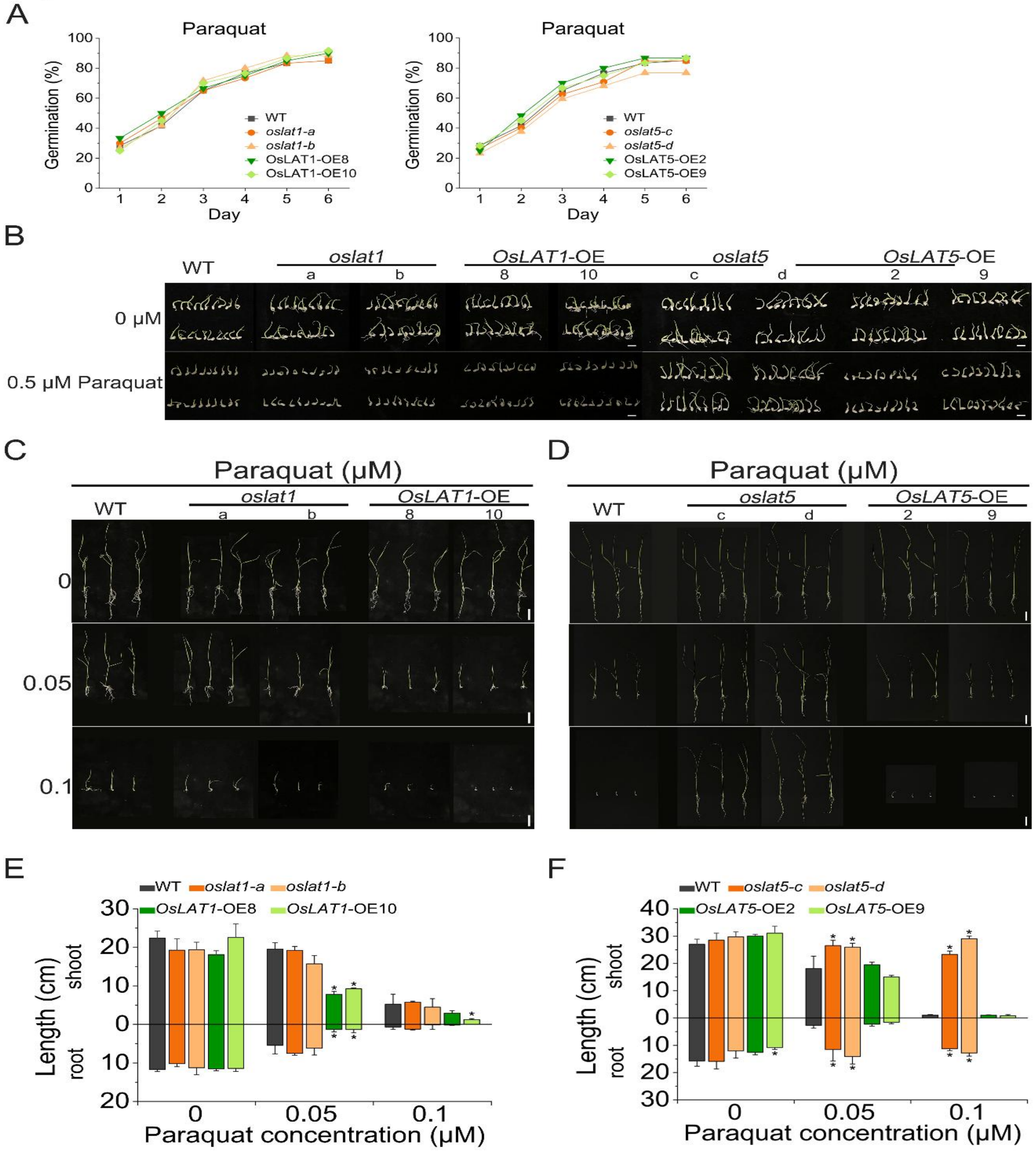


**Fig. S5 Rice seed germination and seedling growth on medium containing paraquat.**

(A) Seed germination curve. Seeds of wild type, mutants and over-expression line of *OsLAT1* or *OsLAT5* were germinated on MS medium added with 0 or 0.5 μM paraquat for 6 days and germination rate was recorded every day.

(B) The above seeds continued to grow followed by six days growing before pictures taken. Bar = 1 cm.

(C-D) Growth of germinated rice seedlings mediated by *OsLAT1* (C) or *OsLAT5* (D) with 0.05 and 0.1 μM paraquat on MS medium. Bar = 4 cm.

(E-F) The shoot and root length of rice seedlings shown in (C) and (D). Data are mean ± SD. n = 3. Asterisks indicate significant differences (one-way ANOVA: *$P < 0.05$).

Figure.S6

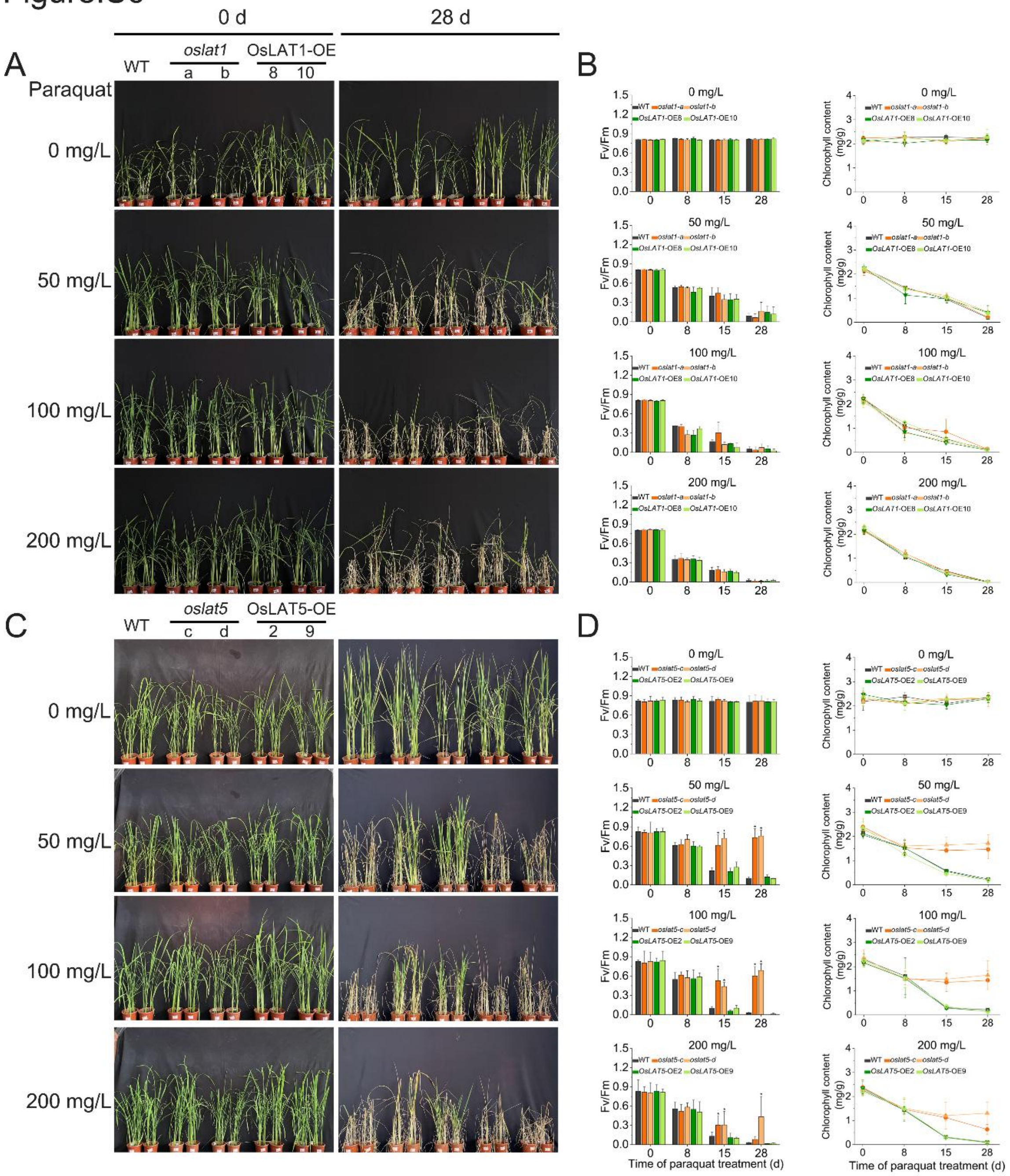


**Fig. S6 Effects of foliar spraying of paraquat on rice growth and development at tillering stage.**

(A) Field-grown rice of different lines of *OsLAT1* (4 weeks old) sprayed with 50, 100 and 200 mg/L paraquat followed by continue growth for different durations.

(B) Analysis of Fv/Fm and chlorophyll contents in the leaves from (A) treated with paraquat for the indicated durations. Data are mean ± SD. n = 3. Asterisks indicate significant differences (one-way ANOVA: $^*P < 0.05$).

(C) Field-grown rice of different lines of *OsLAT5* (4 weeks old) sprayed with 50, 100 and 200 mg/L paraquat followed by continue growth for different durations.

Figure.S7

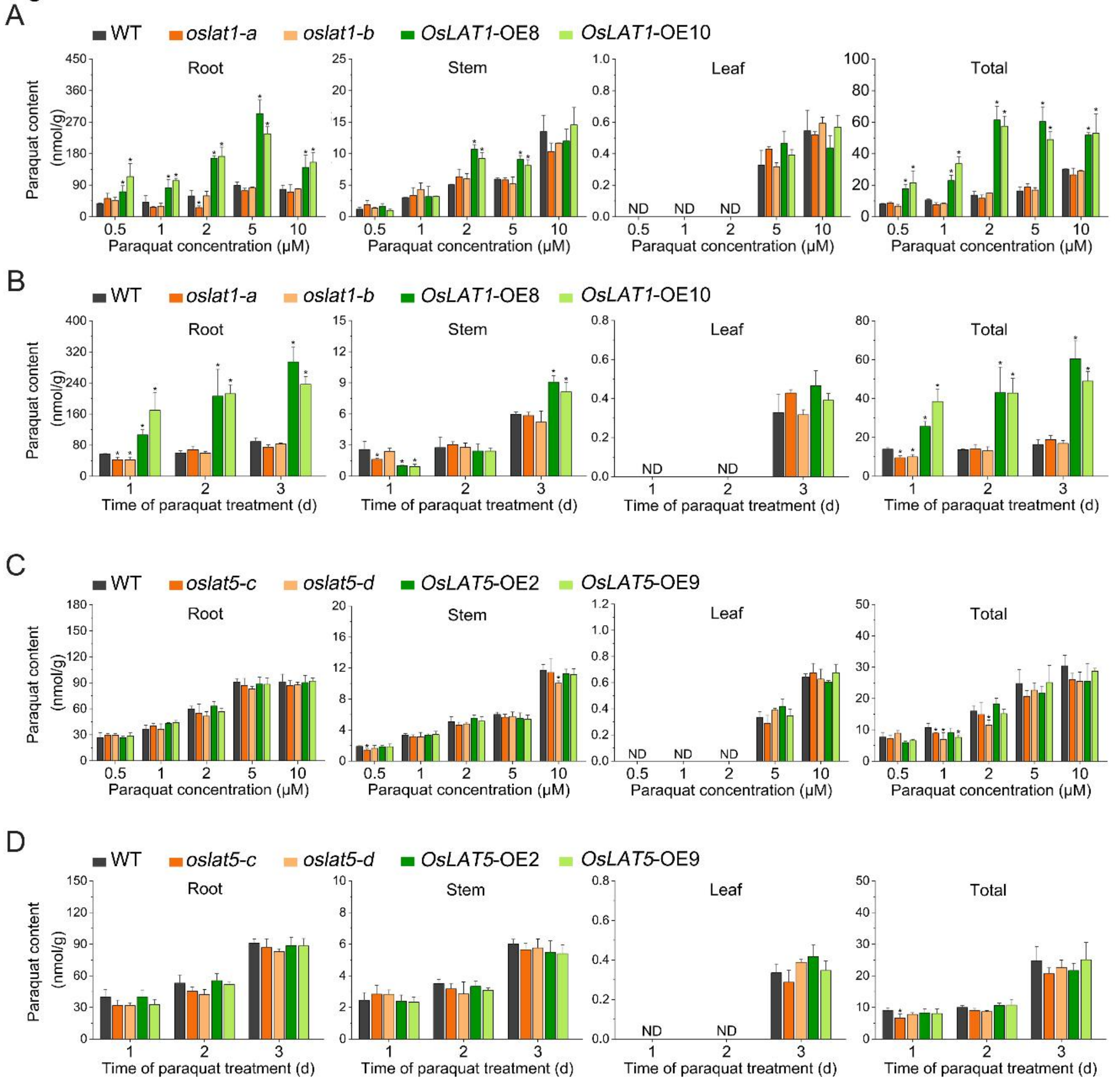


**Fig. S7 Measurement of paraquat uptake in wild type, mutants and overexpressing rice.**

(A and C) Dose-dependent paraquat uptake. The seedlings of different lines of *OsLAT1* (A) or *OsLAT5* (C) (3 weeks old) were incubated with various concentration paraquat for 3 days.

(B and D) Time-dependent paraquat uptake. The seedlings used were as those in (A) or (C), and were incubated with 5 μM paraquat for the time period.

Data are mean ± SD. n = 3. Asterisks indicate significant differences (one-way ANOVA: *$P < 0.05$).

Figure. S8

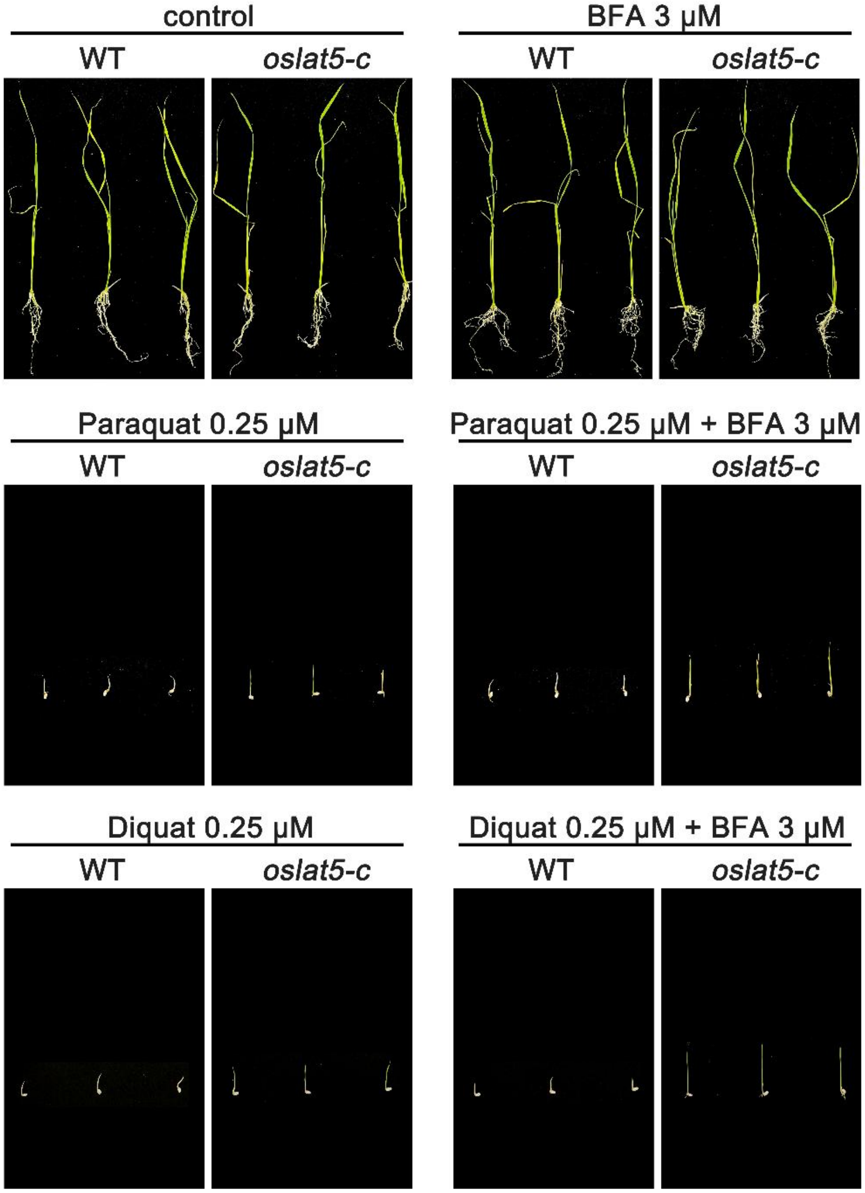


**Fig. S8 Phenotypes of wild type and *oslat5-c* grown on MS plate (control) or MS plate containing 0.25 μM dihydropyridine herbicide and 0.25 μM dihydropyridine herbicide + 3 μM BFA.**

Figure.S9

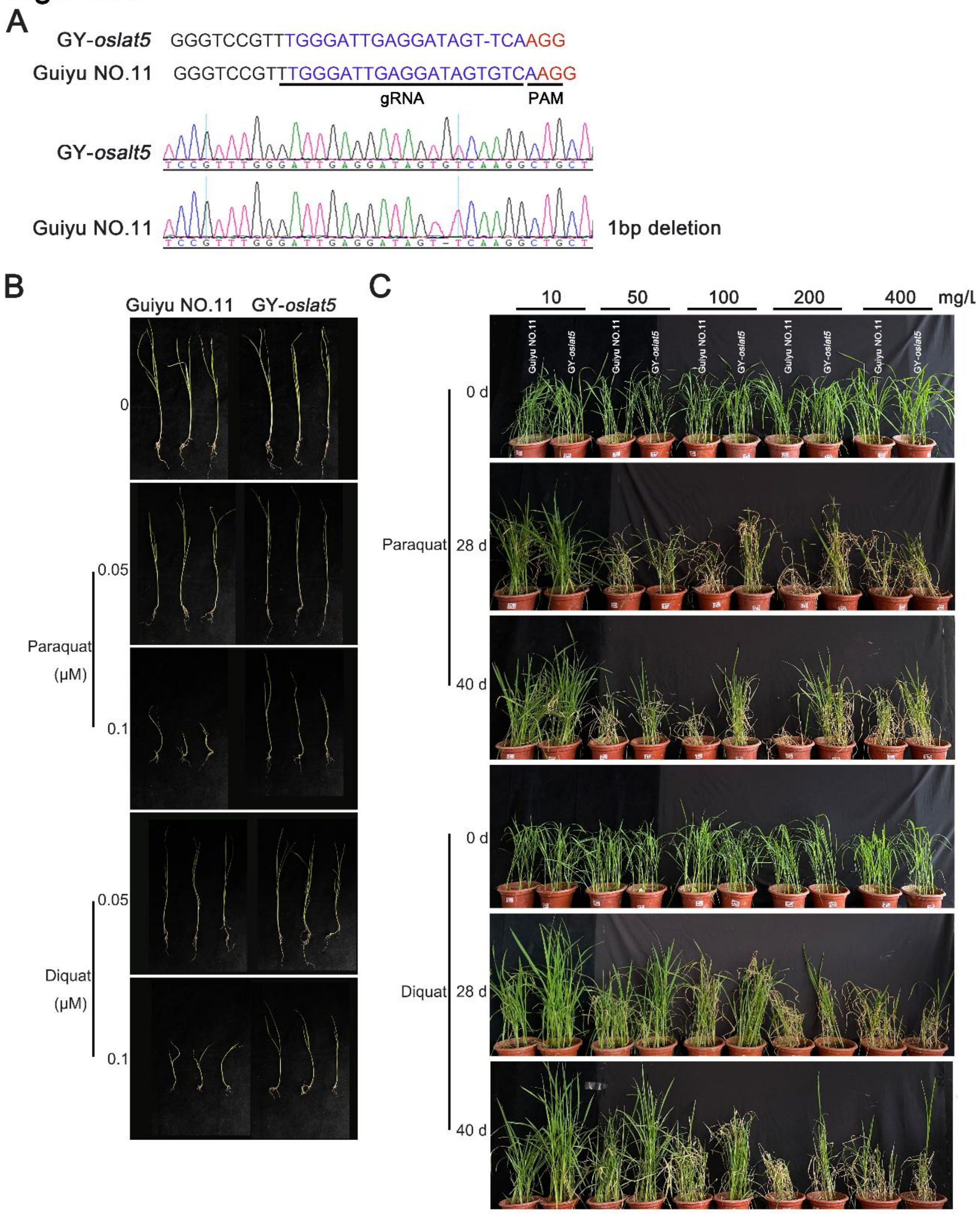


**Fig. S8 The GY-*oslat5* mutant phenotype.**

(A) The *oslat5* knockout mutants (background: Guiyu NO.11) were generated by CRISPR-Cas9 technology, the type of mutation was detected in genomic DNA of GY-*oslat5*.

(B) Effects of dihydropyridine herbicides in rice seed germination of GY-*oslat5*.

(C) Effects of foliar spraying of dihydropyridine herbicides on rice growth and development of GY-*oslat5*.

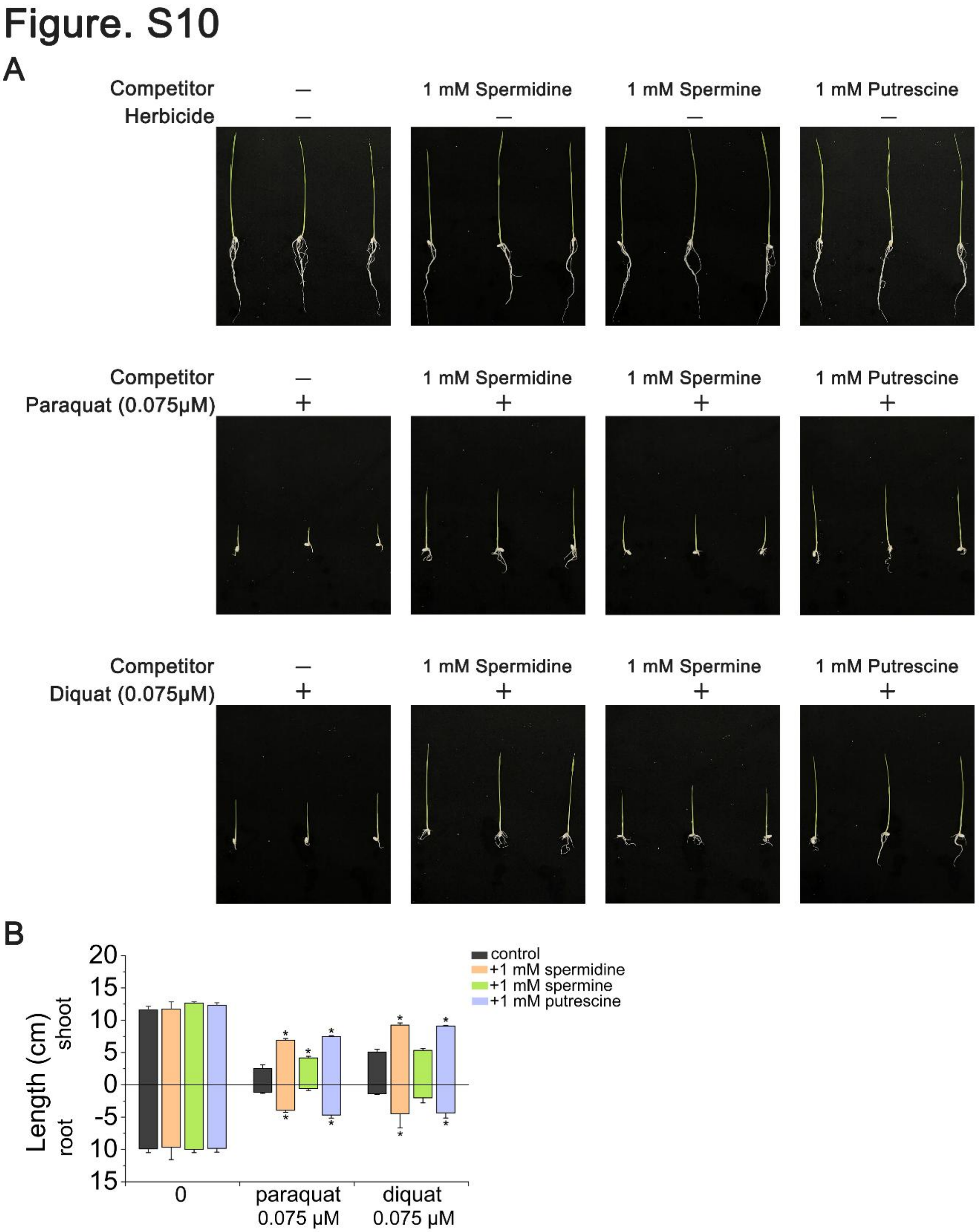


**Fig. S10 Effects of exogenous polyamines on dihydropyridine herbicide toxicity.**

(A) Phenotypes of Zhonghua11 grown on MS plate supplemented with 0.075 μM dihydropyridine herbicide and 1 mM polyamines.

(B) The shoot and root length of rice seedlings shown in (A). Data are mean ± SD. n = 3. Asterisks indicate significant differences (one-way ANOVA: $*P < 0.05$).
